# Proteomics reveals ablation of placental growth factor inhibits the insulin resistance pathways in diabetic mouse retina

**DOI:** 10.1101/338368

**Authors:** Madhu Sudhana Lennikov, Anton Lennikov, Shibo Tang, Hu Huang

## Abstract

The underlying molecular mechanisms that placental growth factor (PlGF) mediates the early complications at non-proliferative diabetic retinopathy (DR) remain largely elusive. The objective of this study is to characterize expression profile due to PlGF ablation in the retina of diabetic mice. The quantitative label-free proteomics was carried out on retinal tissues collected from mouse strains (Akita; PlGF^−/−^ and Akita.PlGF^−/−^). We have identified 3176 total proteins, and 107 were significantly different between the experimental groups, followed by gene ontology, functional pathways, and protein-protein network interaction analysis. Gnb1, Gnb2, Gnb4, Gnai2, Gnao1, Snap25, Stxbp1, Vamp2 and Gngt1 proteins are involved in insulin resistance pathways, which are down-regulated in PlGF ablation in Akita diabetics (Akita.PlGF^−/−^ vs. Akita), up-regulation in Akita vs. C57, PlGF^−/−^ vs. C57. Prdx6, Prdx5 (up-regulation) are known of antioxidant activity; Map2 is involved in neural protection pathways which are up-regulated in Akita.PlGF^−/−^ vs. Akita. Our results suggest that inhibition of insulin resistance pathway and the enhancement of antioxidant defence and neural function may represent the potential mechanisms of anti-PlGF compounds in the treatment of DR.

## 1.0 Introduction

Diabetic retinopathy (DR), a sight-threatening microvascular complication of diabetes myelitis (DM), remains the leading cause of vision loss worldwide in the adult population, especially in economically developed countries. (Lee et al, 2015) With the increasing number of people with DM, the prevalence of DR and diabetic macular edema (DME) is expected to grow. (Wild et al, 2004) Metabolic changes in the diabetic retina result in the altered expression pattern of some mediators including growth factors, neurotrophic factors, cytokines/chemokines, vasoactive agents, and inflammatory and adhesion molecules, resulting in vascular lesions and cell death. (Chen et al, 2015; Kowluru & Mishra, 2015; Liu et al, 2015) Emerging evidence suggests that retinal neurodegeneration is an early event in the pathogenesis of DR which could participate in the development of microvascular abnormalities. (Barber, 2015; Simo et al, 2014) Placenta growth factor (PlGF), a member of VEGF family proteins, first discovered in human placental cDNA in 1991.

In over two decades of scientific research and development have increased our understanding of the PlGF biological function. Despite the high level of expression in placenta, the ablation of PlGF in mice did not compromise the healthy embryonic development or adverse postnatal health effects. (Carmeliet & Jain, 2011) Delivery of recombinant PlGF homodimer, PlGF-VEGFA heterodimer significantly promoted angiogenesis in ischemic conditions through FLT1. (Luttun et al, 2002) Furthermore, many other cell types express PlGF in pathological conditions, including retinal pigment epithelial cells (RPE). (Hollborn et al, 2006) This upregulation is due not only to hypoxia but also from stimulus including nitric oxide (Mohammed et al, 2007), cytokines, as interleukin 1 (IL-1) and TNF-α (De Ceuninck et al, 2004), and transforming growth factor-β1 (TGF-β). (Yao et al, 2005) The observation further confirmed the specific role of PlGF in pathological conditions that during pathological angiogenesis endothelial cells over-express PlGF. (Ponticelli et al, 2008)

Recently our group has reported PlGF deletion in C57BL/6-Ins2Akita/J (Akita) mouse line, containing a dominant mutation that induces spontaneous diabetes with a rapid onset. (Barber et al, 2005) Ablation of PlGF in the diabetic mice resulted in an decreased expression of diabetes-activated hypoxia-inducible factor (HIF)1α, vascular endothelial growth factor (VEGF) pathway, including expression of HIF1α, VEGF, VEGFR1–3, and the extent of phospho (p)-VEGFR1, p-VEGFR2, and p–endothelial nitric oxide synthase, in the retinas of diabetic PlGF^−/−^ mice. Without a noticeable effect on glucose balance or expression of intercellular adhesion molecule-1, vascular cell adhesion molecule-1, CD11b, and CD18 (Huang et al, 2015a).

While many functions and biological roles of PlGF are still currently unknown, the transition to human patients with two phase II clinical trials of anti-PlGF recombinant monoclonal antibody in human DR patients is currently underway. (NCT03071068; NCT03499223; ThromboGenics, 2018) Use of PlGF antibodies in humans presents a challenge of better understanding the functions and pathways involved in PlGF knock out on the proteome scale. Label-free mass spectrometry (LFMS) is a widely used tool for protein identification and quantification it is a gel-free method allowing to conduct whole proteome analysis without the use of isotopic labeling (Luber et al, 2010). Furthermore, as DR affects the expression of many commonly used “housekeeping” proteins such as ACTB and Tubulin, MS approach resolve this issue as proteins are identified by the number of peptide sequences rather than any relative quantification that requires a “housekeeping” protein (Rocha-Martins et al, 2012), (Li & Shen, 2013). In the current study, we used label-free quantitative proteomics analysis to study retinal protein extracts from three genetically modified mouse strains with diabetic and PlGF knockout condition as well as a combination of both to further elucidate the molecular mechanisms of PlGF knockout playing beneficial role in DR on the proteome wide level. We have identified 3176 total proteins, and 107 were significantly different between the experimental groups (p<0.05).

## 1.0 Results

### 2.1 Animals and diabetic conditions

Four 5-6 months old female mice were selected from each strain: C57BL/6-Ins2<Akita>/J.PlGF^−/−^ (Akita.PlGF^−/−^), PlGF^−/−^, C57BL/6-Ins2<Akita>/J (Akita), and C57BL/6J (C57) for this study. Graphical abstract of the animal breeding program is presented in Figure 1A. Animals blood glucose, levels of glycated hemoglobin (HbA1c) and body weight are presented in Table 1. All Akita mice have demonstrated significant increase of blood glucose (BG) levels (p<0.001), HbA1c (p<0.05) and a decrease in body weight (p<0.01) when compared with C57 control animals of the same age. Lack of PlGF did not affect blood glucose (p>0.05) levels of glycated hemoglobin (HbA1c) (p>0.05) and body weight (p>0.05) for Akita.PlGF^−/−^ vs. Akita. There was no significant difference in any of the parameters in PlGF^−/−^ vs. C57 mice of the same age. Retinal protein extracts 4 samples per group were trypsin digested and subjected to the LC/MS/MS analysis (**Figure 1C,D**).

**Table 1:**
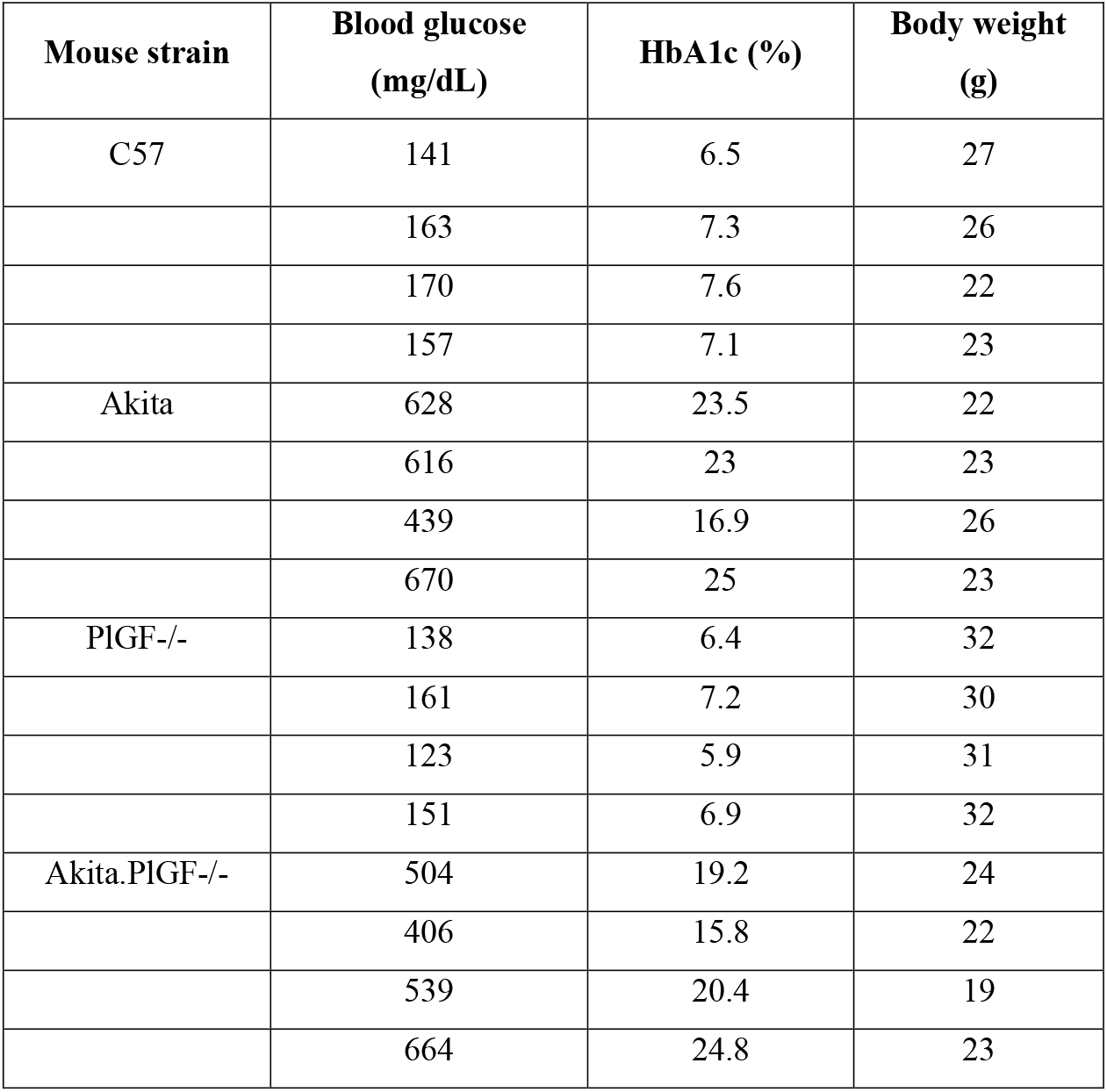
Proteomics samples of blood glucose, levels of glycated hemoglobin (HbA1c) and body weight.

| **Mouse strain** | **Blood glucose**<br>**(mg/dL)** | **HbA1c (%)** | **Body weight**<br>**(g)** |
| --- | --- | --- | --- |
| C57 | 141 | 6.5 | 27 |
|  | 163 | 7.3 | 26 |
|  | 170 | 7.6 | 22 |
|  | 157 | 7.1 | 23 |
| Akita | 628 | 23.5 | 22 |
|  | 616 | 23 | 23 |
|  | 439 | 16.9 | 26 |
|  | 670 | 25 | 23 |
| PIGF-/- | 138 | 6.4 | 32 |
|  | 161 | 7.2 | 30 |
|  | 123 | 5.9 | 31 |
|  | 151 | 6.9 | 32 |
| Akita.PIGF-/- | 504 | 19.2 | 24 |
|  | 406 | 15.8 | 22 |
|  | 539 | 20.4 | 19 |
|  | 664 | 24.8 | 23 |

**Figure 1:**
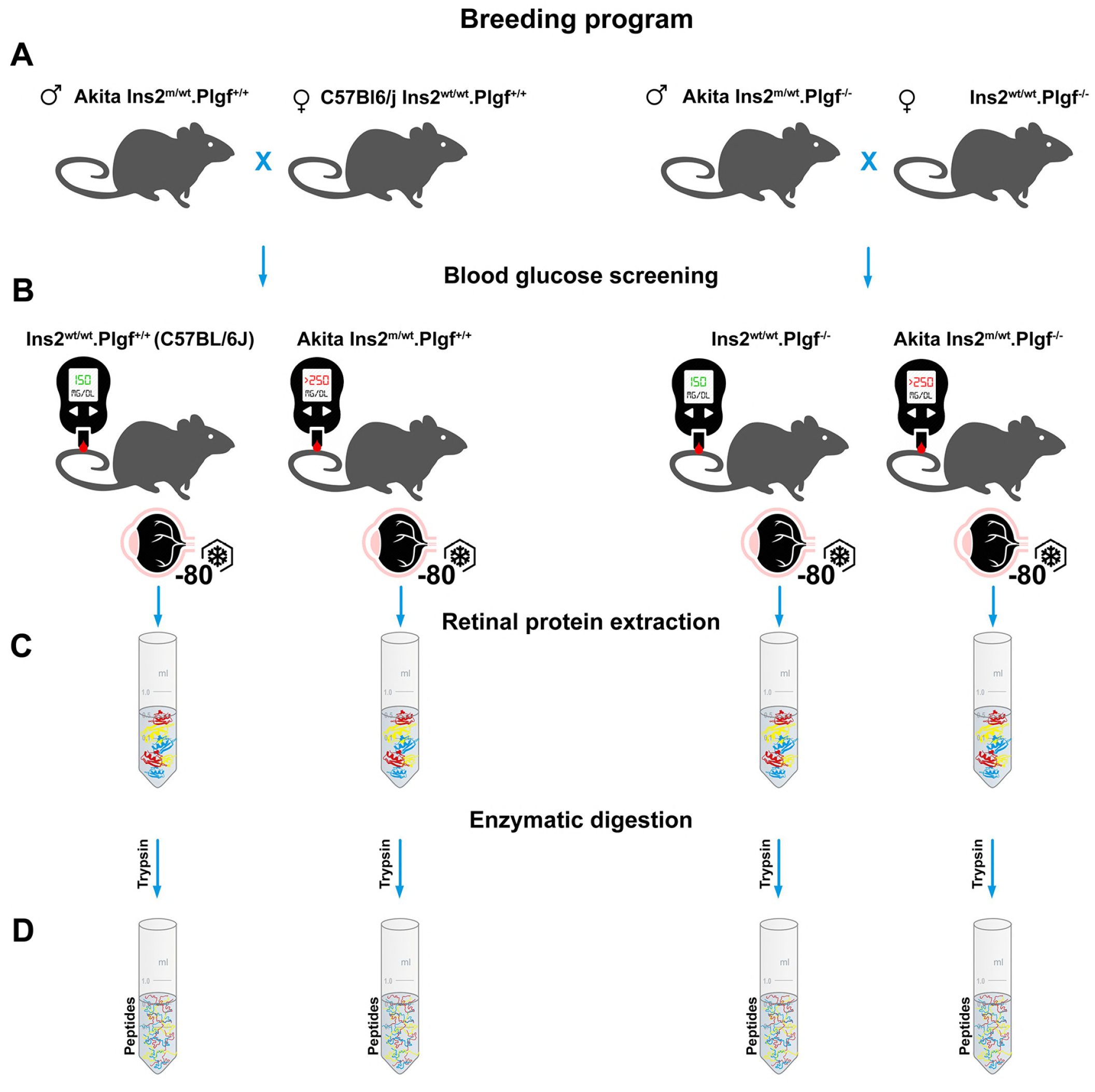
Graphical abstract of the animal breeding program (**A**); screening of diabetic conditions (**B**); retinal sample collection (**C**), initial preparaition of peptide digests (**D**)

### 2.5 Label-free proteomics data analysis

The MaxQuant framework is used for proteomics data analysis, which is written in C# in the Microsoft .NET environment. Algorithmic sets of MaxQuant are freely accessible as source code, and the complete program can be downloaded from www.maxquant.org web link. We followed detailed instructions for installation and support programs Cox et al., 2009. The four experimental data sets (Akita.PlGF^−/−^, PlGF^−/−^, Akita, and C57) were taken as a raw file (**Table 1**) for label-free quantification using MaxQuant version 1.6.01 (http://maxquant.org/). Graphical abstract in **Figure 2** demonstrates the overall analysis workflow. For quantification results of the dataset, MaxQuant calculated the number of quantified proteins, peptides, and sites. The total 200102 msmsScans, 88071 msScans, 5990710 peaks, 465 oxidation(M)Sites, 668368 peptides and 794 proteins were identified (FDR < 0.01) along with mass, m/z, scans, and peaks, etc., for all data sets in the combined/text folder of the MaxQuant output directory.

**Figure 2:**
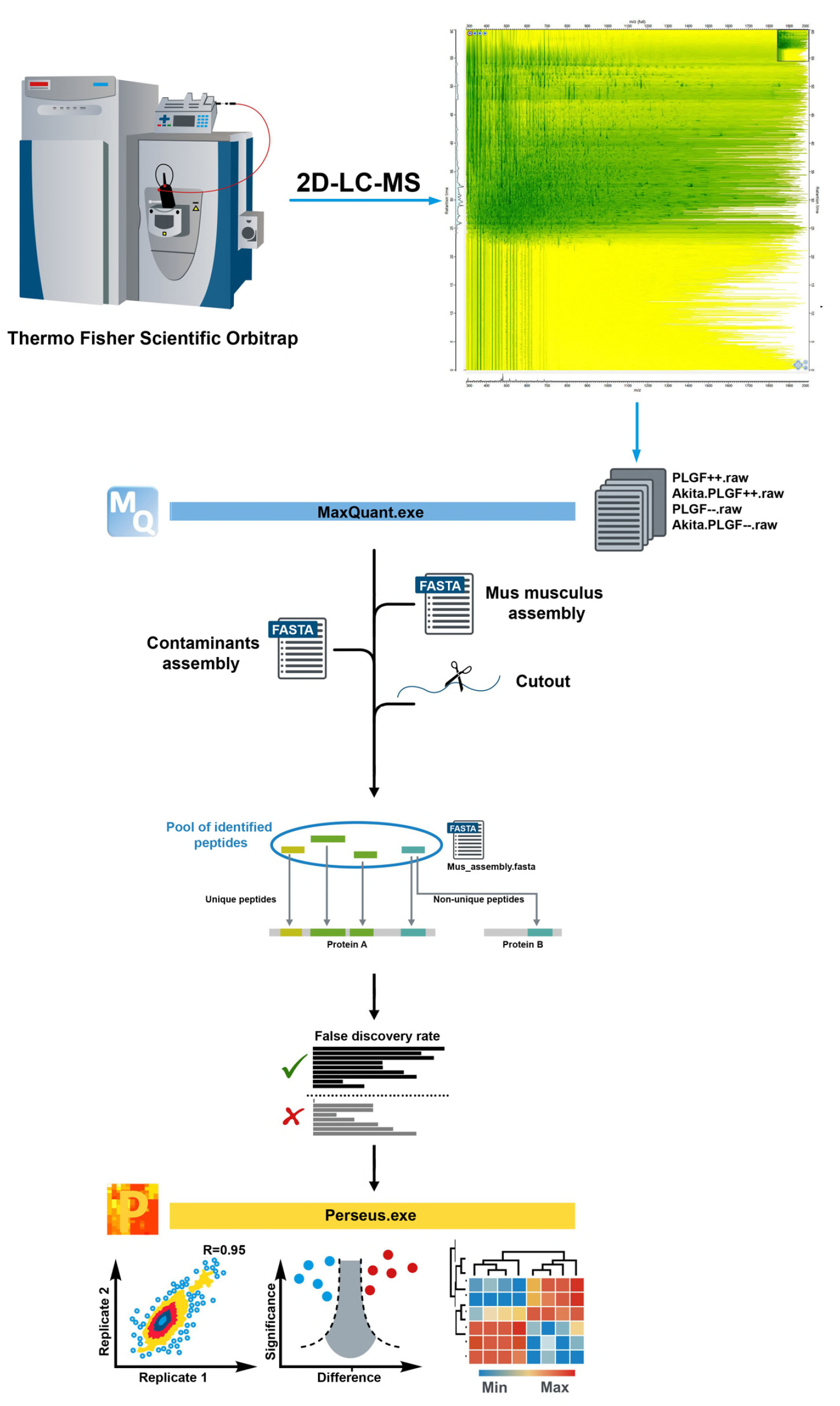
The overall workflow of our proteomics data analysis. The mass spectral (MS) raw data were analyzed with MaxQuant computational proteomics platform (version 1.6.1.0) and its built-in Andromeda search engine. MS data was aligned against Mus musculus UniProtKB (83,598 entries) target database with collection of common contaminants and concatenated with the reversed versions of all sequences. The FDR (false discovery rate) was set to 0.01. Label minimum ratio count was set to 2, peptides for quantification was set to unique and razor and re-quantify to allow identification and quantification of proteins in groups for LFQ analysis. Then for peptide identification, mass and intensity of the peptide peaks in mass spectrometry (MS) spectrum are detected and assembled into three-dimensional (3D) peak hills over the m/z retention time plane. Perseus analysis pipeline, functional classification and pathway analysis.

### 2.6 Perseus analysis pipeline

Perseus is software for shotgun proteomics data analysis, which helps to extract biologically meaningful information from processed raw files. It performs Bioinformatics analyses of the output of MaxQuant, and thus completes the proteomics analysis pipeline. This allows easy integration of an unlimited number of independent statistical tools, which can thus be combined in an analysis. The software already includes various statistical methods and illustrations, such as data transformation, normalization and imputation, unsupervised and supervised learning methods, correlation profiling, enrichment tests, motif identification, volcano plots, scatter plots and more. The MaxQuant software generated output file “proteingroups.txt” was used for statistical analysis by Perseus framework version 1.6.1.1 (Tyanova et al, 2016). The four experimental samples (each one has four individual replicates) were taken as four combinations like Akita.P1GF^−/−^ vs Akita, Akita vs C57, P1GF^−/−^ vs C57 and Akita.P1GF^−/−^ vs P1GF^−/−^ respectively for further statistical analysis. We have estimated the LFQ intensity of all combinations and intensity average is at the same level to the all samples, which describe that the results of LFQ intensity analysis have no biases to the all samples. The correlation coefficient of LFQ intensities was higher than 0.839 in biological replicates. In Akita.P1GF^−/−^ vs Akita combination 0.839 low LFQ intensities between Akita.P1GF^−/−^ replicate 1 and Akita replicate 3, 0.937 high LFQ intensities between Akita.P1GF^−/−^ replicates 1 and 2. (**Suppl. Figure 1B, C, D**) In Akita vs C57 combination 0.867 low LFQ intensities between Akita replicate 4 and C57 replicate 2, 0.943 high LFQ intensities between C57 replicates 3 and 4. (**Suppl. Figure 2B, C, D**) In P1GF^−/−^ vs C57 combination 0.876 low LFQ intensities between P1GF^−/−^ replicate 1 and C57 replicate 3, 0.945 high LFQ intensities between P1GF^−/−^ replicates 2 and 4 (**Suppl. Figure 3B, C, D**). In Akita.P1GF^−/−^ vs P1GF^−/−^ combination 0.877 low LFQ intensities between Akita.P1GF^−/−^ replicate 1 and 3, 0.948 high LFQ intensities between P1GF^−/−^ replicates 2 and 4 (**Suppl. Figure 4B, C, D**), suggesting that the experiment has high repeatability and reliability. The hierarchical clustering and principal component analysis showed clear separation among various components. (**Suppl. Figure 1A, 2A, 3A, 4A**) Based on the threshold like fold-change (FC) of 2 and p-value (p < 0.05) screening differentially expressed proteins (DEPs) in all combinations mentioned above. (**Suppl. Tables 1, 2, 3, 4**). We have also provided MS/MS spectrum and the confidently identified peptide components of Gnb1, Gnb2, Prdx6 and Map2 proteins (**Suppl. Figure 5A, B, C, D**). The Akita.P1GF^−/−^ vs. Akita combination has 31 differentially expressed proteins (6 up-regulate, and 25 downregulate). Prdx6, Map2, Tubb6, Crocc, Hsp90ab1 and Ckb proteins are up-regulated and Gnb2, Snap25, Pcbp1, Atp1a1, Pcbp3, Gnao1, Atp2b1, Atp5o, Gnai2, Vim, Slc25a22, Hnrnpa1, Hist1h1c, Cct8, Hist1h1e, Hist1h1d, Hnrnpc, Atp6v1e1, Hnrnpul2, Mecp2, Hist1h1b, Hnrnpab, Slc6a11, Hnrnpd and Hist1h1a proteins are down-regulated in Akita.P1GF^−/−^ group when compared to Akita. (**Figure 3A**) The Akita vs. C57 combination has 26 DEPs, 15 proteins are up-regulated, and 11 proteins are down-regulated. Hnrnpd, Hnrnpc, Hnrnpab, Atp1b2, Hnrnpa1, Lmnb2, Atp5o, Gnb1, Pcbp1, Gnb2, Atp6v1a, Atp1a2, Atp1a1, Hnrnpk and Stxbp1 proteins are up-regulated and Glul, Cfl1, Prdx6, Hba2, Map2, Epb41l3, Ntm, Uqcrq, Gfap, Nme1 and Nefm proteins are down-regulated in Akita group when compared to C57 (**Figure 4A**). The P1GF^−/−^ vs. C57 combination has 31 DEPs, 15 proteins are up-regulated, and 16 proteins are down-regulated. Ndufa8, Cirbp, Rbp3, Napb, Crocc, Gngt1, Gnat1, Atp1b2, Gnb1, Atp5o, Lmnb2, Atp5f1, Gnb4, Gnb2 and Aco2 proteins are up-regulated and Nono, Tubb3, Vamp1, Ywhab, Srsf1, Hnrnpc, Hba2, Ywhag, Epb4.1l3, Tpm1, Eif4a1, Rac1, Gfap, Hnrnpa0, Nefm and Rhobtb3 are down-regulated proteins in P1GF^−/−^ group when compared to C57 (**Figure 5A**). The Akita.P1GF^−/−^ vs. P1GF^−/−^ combination has 19 DEPs, 6 proteins are up-regulated, and 13 proteins are down-regulated. Tubb6, Tubb3, Hsp90ab1, Tubb2b, Tubb5, and Hspa8 are up-regulated Atp5o, Map1b, Immt, Hnrnpm, Vdac2, Pura, Cct8, Cirbp, Cox5b, Mecp2, Rs1, Ndufa8 and Erh proteins are down-regulated in Akita.P1GF^−/−^ group when compared to P1GF^−/−^ (**Figure 6A**). In all combinations of DEPs are used for gene annotation (GO) and pathways analysis.

**Figure 3:**
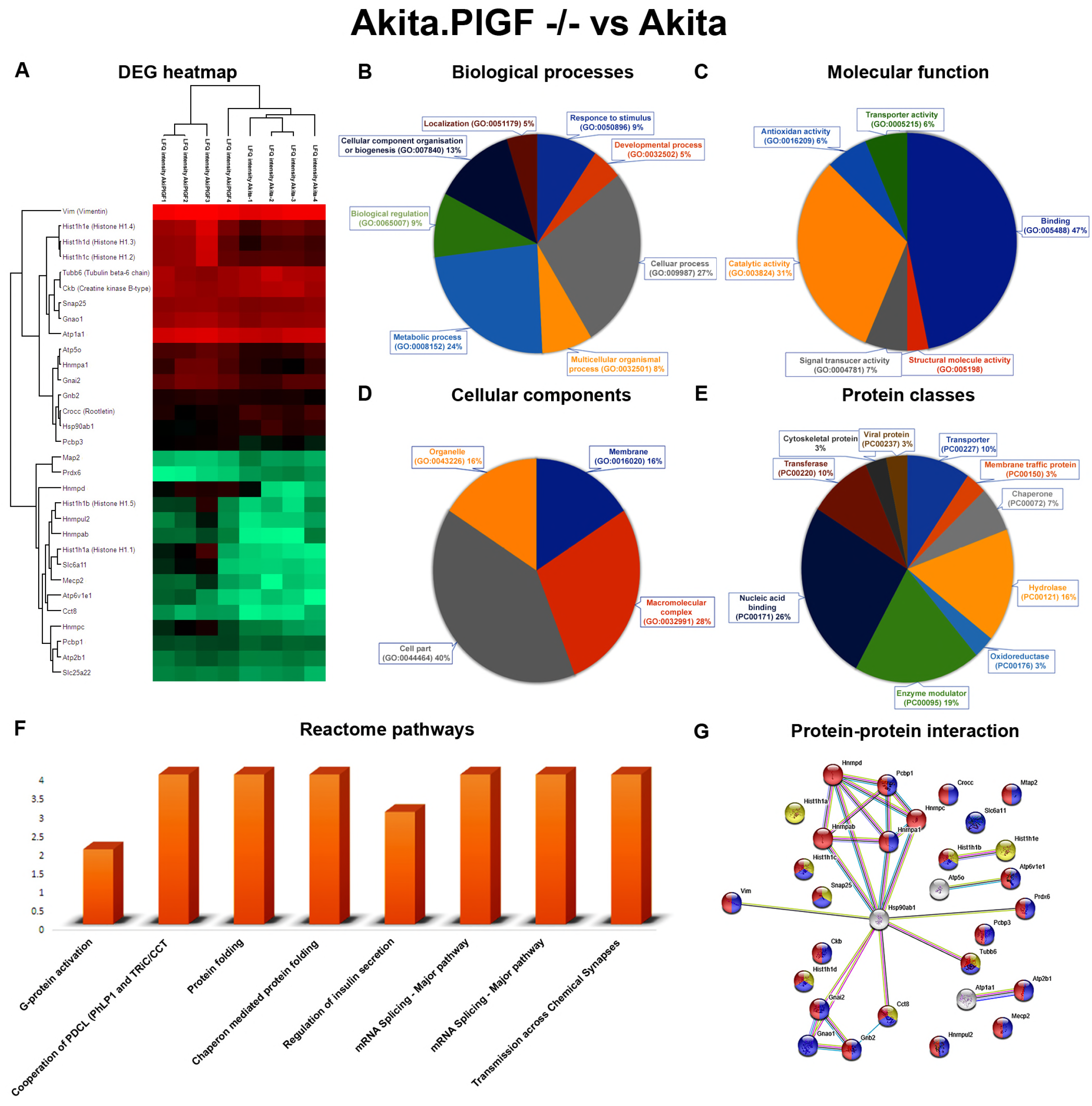
Functional annotation, reactome pathways and protein-protein interaction network of Akita.P1GF−/−vs Akita. (**A**) Significantly found up and down-regulated proteins were represented as a heatmap. Green and red colour indicates low to high intensity of abundance of proteins. (**B**) The differentially expressed proteins are involved in different biological processes like biological regulation (GO:0065007), cellular component organization or biogenesis (GO:0071840), cellular process (GO:0009987), localization (GO:0051179), metabolic process (GO:0008152), multicellular organismal process (GO:0032501) and response to stimulus (GO:0050896) respectively. (**C**) The differentially expressed proteins are involved in different molecular functions like antioxidant activity (GO:0016209), binding (GO:0005488), catalytic activity (GO:0003824), signal transducer activity (GO:0004871), structural molecule activity (GO:0005198) and transporter activity (GO:0005215) respectively. (**D**) The differentially expressed proteins are involved in various cellular components functions like cell part (GO:0044464), macromolecular complex (GO:0032991), membrane (GO:0016020), and organelle (GO:0043226) respectively. (**E**) The differentially expressed proteins are different classes of proteins like chaperone, cytoskeletal protein, enzyme modulator, hydrolase, membrane traffic protein, nucleic acid binding, oxidoreductase, transferase, transporter and viral proteins respectively. (**F**) The differentially expressed proteins are involved in various reactome biological pathways like G-protein activation pathway, Co-operation of PDCL (PhLP1) and TRiC/CCT in G-protein beta folding pathway, Chaperonin-mediated protein folding pathway, Protein folding pathway, Regulation of insulin secretion pathway, mRNA Splicing - Major Pathway and Transmission across Chemical Synapses pathway respectively. (**G**) Highlighted the various colours of up and down-regulated proteins involved in complex protein assembly (yellow) in biological process (GO:0006461), binding (blue) function of molecular function (GO:0005488), membrane-bounded organelle (red) proteins in cellular component (GO:0043227) respectively.

**Figure 4:**
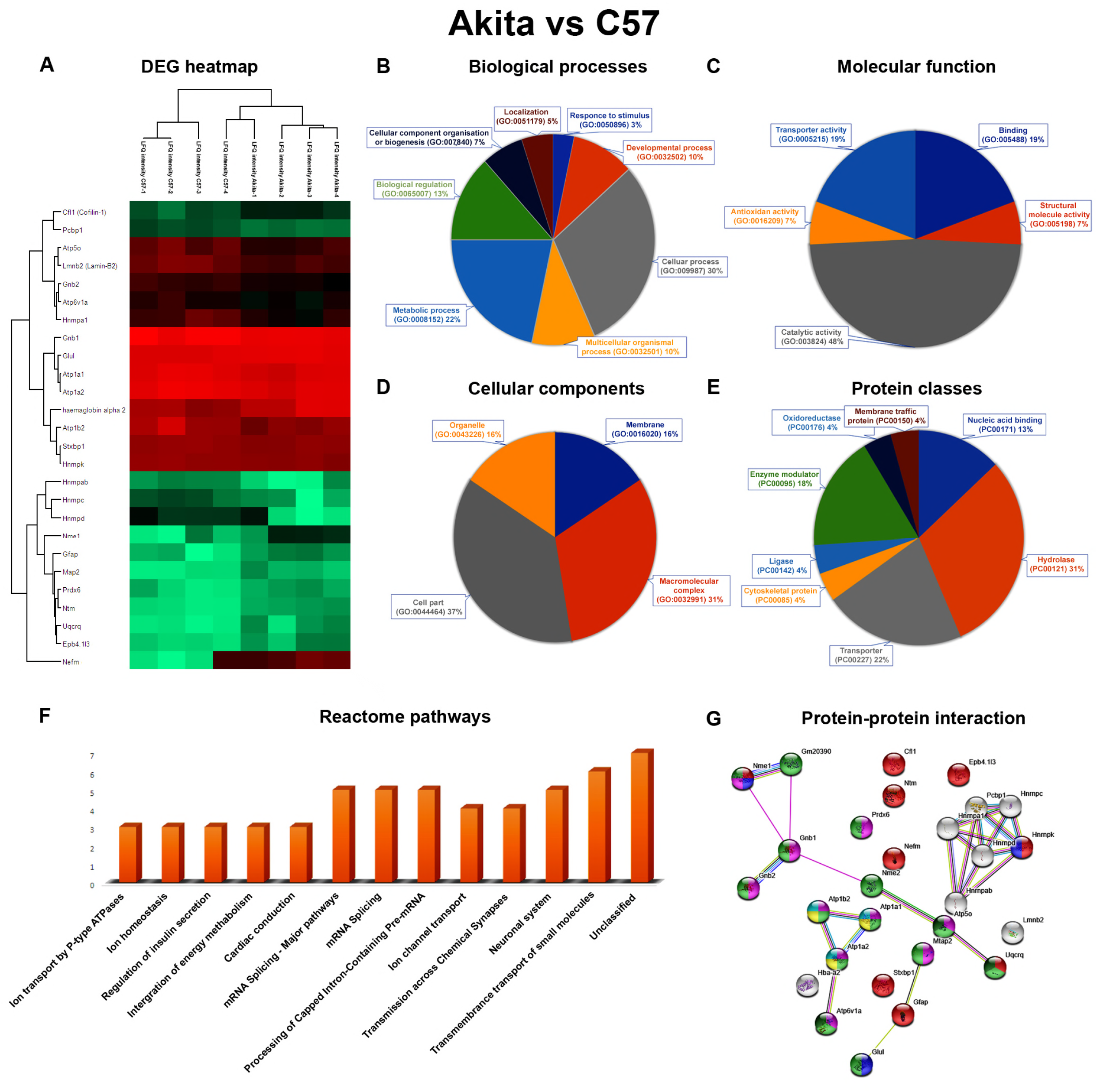
Functional annotation, reactome pathways and protein-protein interaction network of Akita vs C57. (**A**) Significantly found up and down-regulated proteins were represented as a heatmap. Green and red colour indicates low to high intensity of abundance of proteins (**B**) The differentially expressed proteins are involved in different biological processes such as biological regulation (GO:0065007), cellular component organization or biogenesis (GO:0071840), cellular process (GO:0009987), developmental process (GO:0032502), localization (GO:0051179), metabolic process (GO:0008152), multicellular organismal process (GO:0032501), response to stimulus (GO:0050896) respectively. (**C**) The differentially expressed proteins are involved in various molecular functions such as antioxidant activity (GO:0016209), binding (GO:0005488), catalytic activity (GO:0003824), structural molecule activity (GO:0005198) and transporter activity (GO:0005215) respectively. (**D**) The differentially expressed proteins are involved in various cellular components functions like cell part (GO:0044464), macromolecular complex (GO:0032991), membrane (GO:0016020) and organelle (GO: 0043226) respectively. (**E**) The differentially expressed proteins are different classes of proteins like cytoskeletal protein, enzyme modulator, hydrolase, ligase, membrane traffic protein, nucleic acid binding, oxidoreductase and transporter respectively. (**F**) The differentially expressed proteins are involved in various reactome biological pathways like Ion transport by P-type, Ion channel transport, Transmembrane transport of small molecules, Ion homeostasis (Atp1a1, Atp1b2 and Atp1a2), Cardiac conduction, Regulation of insulin secretion, Integration of energy metabolism, mRNA Splicing - Major Pathway, Processing of Capped Intron-Containing Pre-mRNA, Transmission across Chemical Synapses, Neuronal System and Unclassified pathways respectively. (**G**) Highlighted the various colours of up and down-regulated proteins involved in nervous system development (GO:0007399) (red), response to glucose (GO:0009749) (blue) of biological process, catalytic activity (GO:0003824) (light green), hydrolase activity (GO:0016787) (pink), of molecular functions, Insulin secretion (ko04911) (yellow), Pancreatic secretion (ko04972) (cyan), Oxidative phosphorylation (ko00190) (thick green) of KEGG pathways.

**Figure 5:**
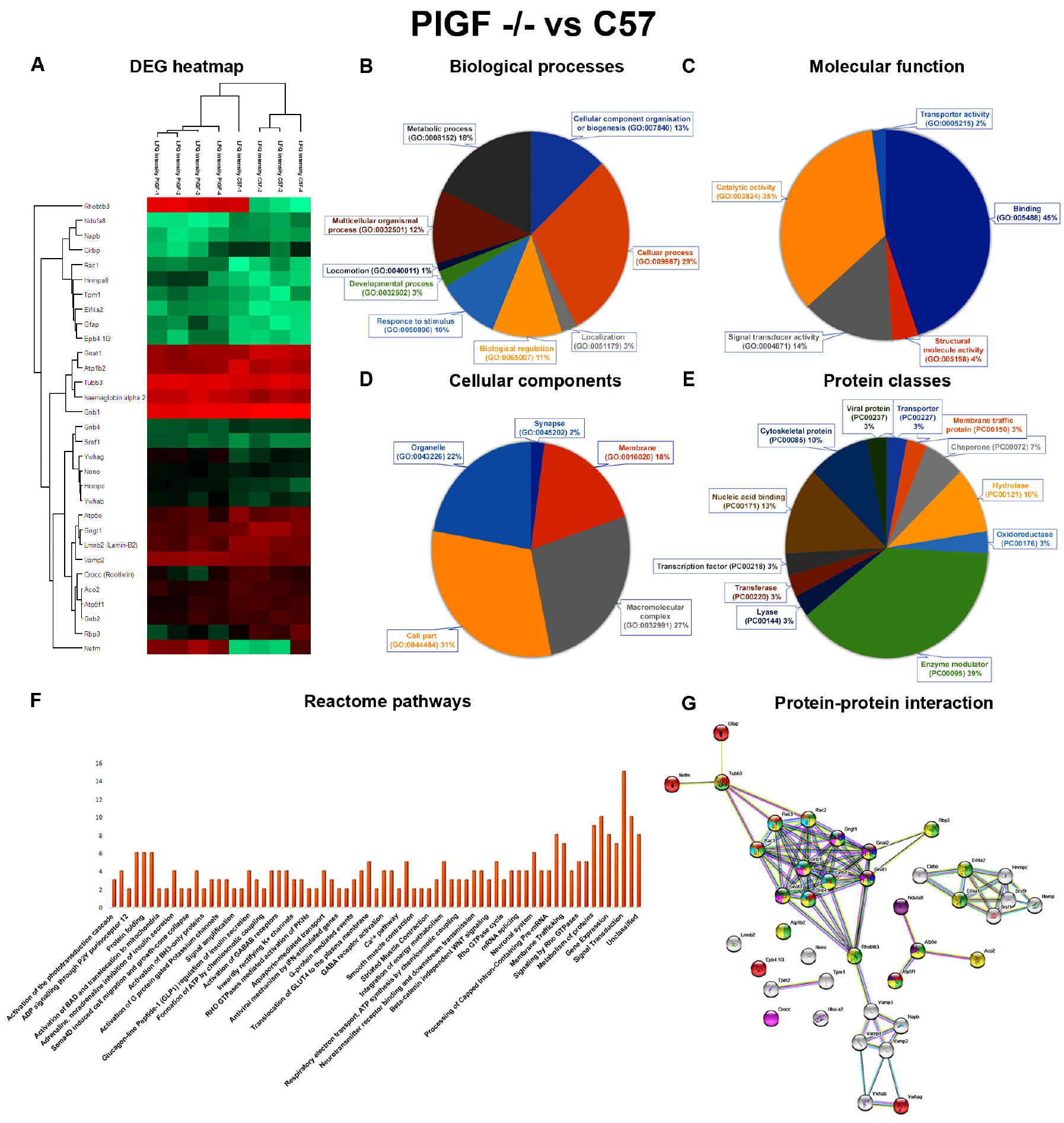
Functional annotation, reactome pathways and protein-protein interaction network of P1GF^−/−^ vs C57. (**A**) Significantly found up and down-regulated proteins were represented as a heatmap. Green and red colour indicates low to high intensity of abundance of proteins (**B**) The differentially expressed proteins are involved in different biological processes such as biological regulation (GO:0065007), cellular component organization or biogenesis (GO:0071840), cellular process (GO:0009987), developmental process (GO:0032502), localization (GO:0051179), locomotion (GO:0040011), metabolic process (GO:0008152), multicellular organismal process (GO:0032501), response to stimulus (GO:0050896) respectively. (**C**) The differentially expressed proteins are involved in various molecular functions such as binding (GO:0005488), catalytic activity (GO:0003824), signal transducer activity (GO:0004871), structural molecule activity (GO:0005198), transporter activity (GO:0005215) respectively. (**D**) The differentially expressed proteins are involved in various cellular components functions like cell part (GO:0044464), macromolecular complex (GO:0032991), membrane (GO:0016020), organelle (GO:0043226), and synapse (GO:0045202) respectively. (**E**) The differentially expressed proteins are classified in to various protein class such as chaperone, cytoskeletal protein, enzyme modulator, hydrolase, membrane traffic protein, nucleic acid binding, oxidoreductase, transcription factor, transferase, transporter, and viral protein respectively. (**F**) The differentially expressed proteins are involved in various reactome biological pathways like Activation of the phototransduction cascade, Signal Transduction, G-protein activation, Opioid Signalling, Signalling by GPCR, ADP signalling through P2Y purinoceptor-12, Platelet activation, signalling and aggregation, Hemostasis, Cooperation of PDCL (PhLP1) and TRiC/CCT in G-protein beta folding, Chaperonin-mediated protein folding, Metabolism of proteins, Activation of BAD and translocation to mitochondria, Intrinsic Pathway for Apoptosis, Adrenaline, noradrenaline inhibits insulin secretion, Regulation of insulin secretion, Metabolism, RHO GTPase Effectors, Signaling by Rho GTPases, Glucagon-type ligand receptors, Activation of G protein gated Potassium channels, Transmission across Chemical Synapses, Glucagon-like Peptide-1 (GLP1) regulates insulin secretion, Inhibition of voltage gated Ca2+ channels via Gbeta/gamma subunits, Neuronal System, Inactivation, recovery and regulation of the phototransduction cascade, Beta-catenin independent WNT signalling, Rho GTPase cycle and un classified pathway respectively. (**G**) Highlighted the various colours of up and down-regulated proteins involved in nervous system development (GO:0007399) (red), eye photoreceptor cell differentiation (GO:0001754) (blue) of biological process, hydrolase activity (GO:0016787) (light green), catalytic activity (GO:0003824) (yellow) of molecular functions, photoreceptor inner segment (GO:0001917) (pink), photoreceptor outer segment (GO:0001750) (thick green) of cellular components, Ras signalling pathway (ko04014) (cyan), VEGF signalling pathway (ko04370) (orange), Oxidative phosphorylation (ko00190) (magenta) of KEGG pathways.

**Figure 6:**
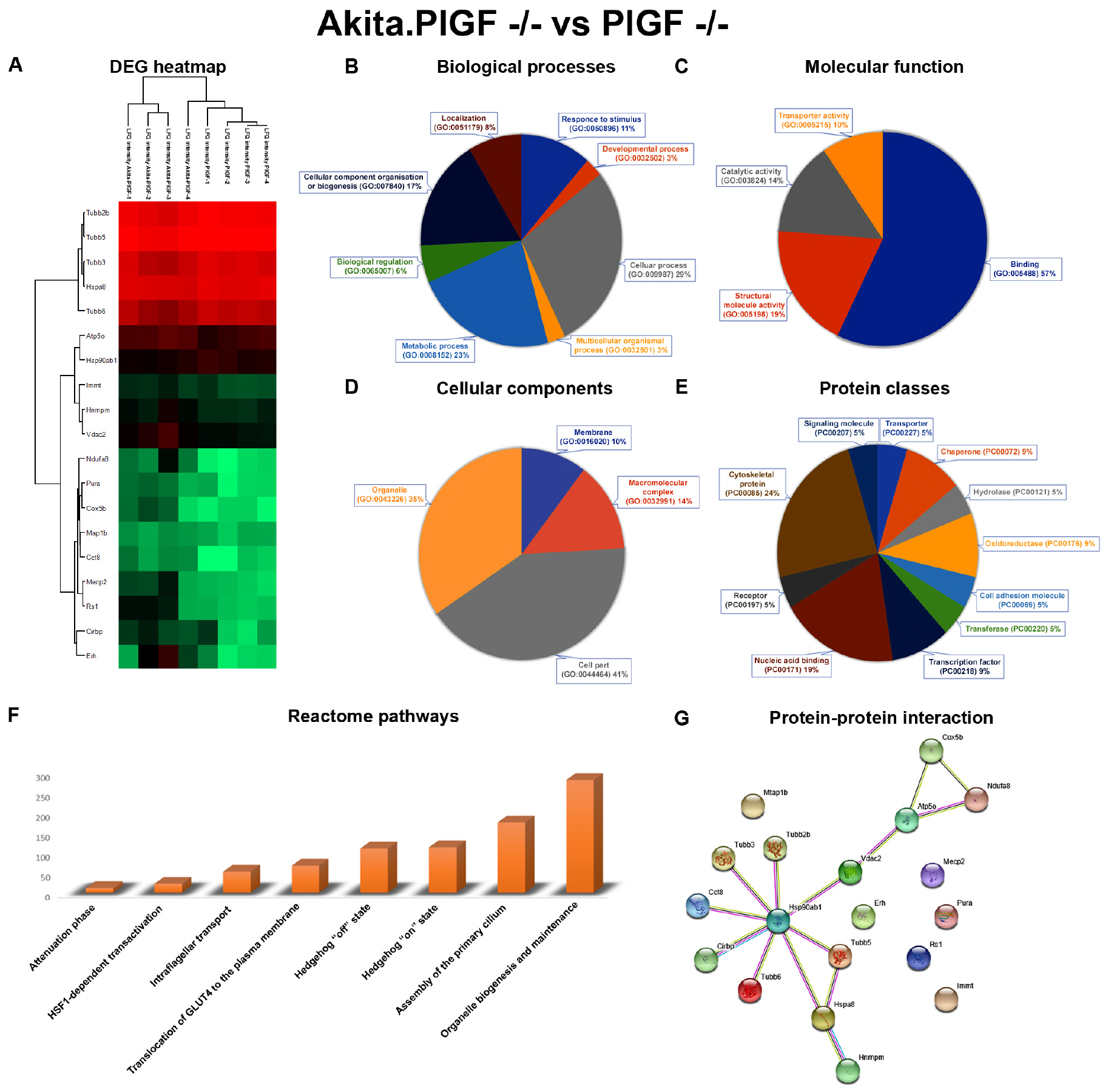
Functional annotation, reactome pathways and protein-protein interaction network of Akita.P1GF^−/−^ vs P1GF^−/−^. (**A**) Significantly up and down-regulated proteins were represented as a heatmap. Green and red colour indicates low to high intensity of abundance of proteins. (**B**) The differentially expressed proteins are involved in different biological processes such as biological regulation (GO:0065007), cellular component organization or biogenesis (GO:0071840), cellular process (GO:0009987), developmental process (GO:0032502), localization (GO:0051179), metabolic process (GO:0008152), multicellular organismal process (GO:0032501), and response to stimulus (GO:0050896) respectively. (**C**) The differentially expressed proteins are involved in various molecular functions such as binding (GO:0005488), catalytic activity (GO:0003824), structural molecule activity (GO:0005198), and transporter activity (GO:0005215) respectively. (**D**) The differentially expressed proteins are involved in various cellular components functions like cell part (GO:0044464), macromolecular complex (GO:0032991), membrane (GO:0016020) and organelle (GO:0043226) respectively (**E**) The differentially expressed proteins are classified in to various protein class such as cell adhesion molecule, chaperone, cytoskeletal protein, hydrolase, nucleic acid binding, oxidoreductase, receptor, signalling molecule, transcription factor, transferase and transporter respectively. (**F**) The differentially expressed proteins are involved in various reactome biological pathways like Attenuation phase, HSF1-dependent transactivation, Intraflagellar transport, Assembly of the primary cilium, Organelle biogenesis and maintenance, Translocation of GLUT4 to the plasma membrane, Hedgehog ‘off’ state, and Hedgehog ‘on’ state respectively. (**G**). Highlighted the various colours of up and down-regulated proteins involved in protein folding (GO:0006457) (blue) of biological process, binding (GO:0005488) (red) of molecular function, intracellular organelle part (GO:0044446) (green) of cellular component, Gap junction (ko04540) (green) of KEGG pathway.

### 2.7 Functional classification and pathway analysis

The functional analysis is a crucial factor step in data analysis for no functional annotation of the data sets. The DEPs were connecting to at least one annotation term each within the molecular function (MF), biological process (BP) and cellular component (CC) classes. In this study, DAVID Bioinformatics Resources 6.8 (https://david.ncifcrf.gov/) and GO Enrichment Analysis (http://geneontology.org/page/go-enrichment-analysis) are used for gene annotation of DEPs. All the up and down regulated individual combinations of DEPs were uploaded to the DAVID annotation tool using complete mouse proteome as background. The GO terms were predicted based on Expression Analysis Systematic Explorer (EASE) < 0.1 and threshold count (TC) ≥ 2. The molecular functions, biological process, cellular components protein classes, and pathways were predicted in the significant enriched GO terms of up and down-regulated proteins.

The Akita.P1GF^−/−^ vs P1GF^−/−^ combination DEPs were involved in different biological processes like biological regulation (GO:0065007), cellular component organization or biogenesis (GO:0071840), cellular process (GO:0009987), localization (GO:0051179), metabolic process (GO:0008152), multicellular organismal process (GO:0032501) and response to stimulus (GO:0050896) respectively (**Figure 2B**). Gnao1, Gnai2, Atp2b1, Atp1a1, Snap25 and Slc6a11 are involved in biological regulation (GO:0065007), Hist1h1d, Tubb6, Cct8, Map2, Hist1h1c, Hist1h1a, Snap25, Hist1h1b and Hist1h1e are involved in cellular component organization or biogenesis (GO:0071840), Slc25a22, Gnao1, Atp2b1, Pcbp1, Hist1h1d, Gnai2, Tubb6, Cct8, Map2, Prdx6, Gnb2, Hist1h1c, Hist1h1a, Pcbp3, Snap25, Hist1h1b, Atp1a1, Slc6a11, Crocc and Hist1h1e are involved in cellular process (GO:0009987), Pcbp1, Map2, Pcbp3 are involved in localization (GO:0051179), Slc25a22, Gnao1, Hsp90ab1, Atp2b1, Pcbp1, Hist1h1d, Gnai2, Cct8, Ckb, Prdx6, Hist1h1c, Hist1h1a, Pcbp3, Hist1h1b, Atp1a1 and Hist1h1e are involved in metabolic process (GO:0008152), Pcbp1, Map2, Gnb2, Pcbp3, Snap25 and Slc6a11 are involved in multicellular organismal process (GO:0032501), Gnao1, Hsp90ab1, Gnai2, Mecp2, and Prdx6 are involved in response to stimulus (GO:0050896) respectively. The DEPs were involved in different molecular functions like antioxidant activity (GO:0016209), binding (GO:0005488), catalytic activity (GO:0003824), signal transducer activity (GO:0004871), structural molecule activity (GO:0005198) and transporter activity (GO:0005215) (**Figure 2C**). Prdx6, Prdx5 have antioxidant activity, Gnao1, Atp2b1, Pcbp1, Hist1h1d, Gnai2, Mecp2, Tubb6, Cct8, Map2, Gnb2, Hist1h1c, Hist1h1a, Pcbp3, Snap25, Hist1h1b, Crocc, and Hist1h1e have binding activity, Gnao1, Atp2b1, Pcbp1, Gnai2, Ckb, Prdx6, Gnb2, Ckb, Pcbp3, Atp1a1 and Crocc have catalytic activity, Gnao1 and Gnai2 have signal transducer activity, Tubb6 has structural molecule activity and Slc25a22, Atp2b1, Atp1a1 and Slc6a11 have transporter activity respectively. The DEPs were involved in various cellular components functions like cell part (GO:0044464) (Gnao1, Atp2b1, Gnai2, Mecp2, Tubb6, Cct8, Map2, Prdx6, Snap25 and Slc6a11), macromolecular complex (GO:0032991) (Gnao1, Pcbp1, Gnai2, Cct8, Gnb2, Pcbp3 and Snap25), membrane (GO:0016020) (Gnao1, Atp2b1, Gnai2 and Snap25) and organelle (GO:0043226) (Atp2b1, Mecp2, Tubb6 and Cct8) respectively (**Figure 2D**). DEPs are belonging to different classes of proteins like chaperone (Hsp90ab1 and Cct8), cytoskeletal protein (Tubb6), enzyme modulator (Gnao1, Pcbp1, Gnai2, Gnb2, Pcbp3 and Crocc), hydrolase (Atp2b1, Pcbp1, Gnb2, Pcbp3 and Atp1a1), membrane traffic protein (Snap25), nucleic acid binding (Pcbp1, Hist1h1d, Mecp2, Hist1h1c, Hist1h1a, Pcbp3, Hist1h1b and Hist1h1e), oxidoreductase (Prdx6), transferase (Mecp2 and Ckb), transporter (Atp2b1, Atp1a1 and Slc6a11) and viral protein (Crocc) respectively (Figure 2E). The up and down regulated proteins were involved in various reactome biological pathways like G-protein activation pathway (Gnao1 and Gnai2), Co-operation of PDCL (PhLP1) and TRiC/CCT in G-protein beta folding pathway (Gnao1, Gnai2, Cct8 and Gnb2), Chaperonin-mediated protein folding pathway (Gnao1, Gnai2, Cct8 and Gnb2), Protein folding pathway (Gnao1, Gnai2, Cct8 and Gnb2), Regulation of insulin secretion pathway (Gnai2, Gnb2 and Snap25), mRNA Splicing - Major Pathway (Hnrnpa1, Pcbp1, Hnrnpc and Hnrnpd), Transmission across Chemical Synapses pathway (Gnai2, Gnb2, Snap25 and Slc6a11) respectively (**Figure 2F**). **Table 2** showed the up and down-regulated proteins, gene name, MS/MS count, unique sequence coverage, molecular weight (MW), T-test difference, p-value, various KEGG name, and pathways.

**Table 2:** The list of significantly differentially expressed proteins, gene name, MS/MS count, unique sequence coverage, molecular weight (MW), T-test difference, p-value, various KEGG name, and pathways in Akita.PLGF^−/−^ group compared to Akita group.

| Protein names | Gene names | Unique sequence coverage [%] | Mol. weight [kDa] | MS/MS count | Student's T-test Difference AkiPLGF<sup>-/-</sup>_Akc57 | Student's T-test Test statistic AkiPLGF<sup>-/-</sup>_Akc57 | P-value | KEGG | KEGG name |
| --- | --- | --- | --- | --- | --- | --- | --- | --- | --- |
| Peroxiredoxin-6 | Prdx6 | 22.3 | 24.826 | 25 | 1.82912302 | 5.092553817 | 2.6501 | ko00360<br>ko00680<br>ko00940 | Methane metabolism;<br>Phenylalanine metabolism;<br>Phenylpropanoid biosynthesis |
| Microtubule-associated protein;<br>Microtubule-associated protein 2 | Map2 | 9.7 | 49.264 | 46 | 0.723612309 | 2.727109127 | 1.4644 | ko04012<br>ko04722<br>ko04728<br>ko04310 | ErbB signaling pathway<br>Neurotrophin signaling pathway<br>Dopaminergic synapse<br>Wnt signaling pathway |
| Tubulin beta-6 chain | Tubb6 | 4 | 50.09 | 174 | 0.552398205 | 2.556084196 | 1.3651 | ko04145<br>ko04540<br>ko05130 | Gap junction<br>Pathogenic Escherichia coli infection<br>Phagosome |
| Rootletin | Crocc | 18.3 | 226.94 | 143 | 0.509349823 | 4.191491283 | 2.2411 |  |  |
| Heat shock protein HSP 90-beta | Hsp90ab1 | 8.3 | 83.28 | 83 | 0.483288288 | 3.900802892 | 2.0981 | ko04141<br>ko04612<br>ko04621<br>ko04626<br>ko04914<br>ko05200<br>ko05215 | Antigen processing and presentation<br>NOD-like receptor signaling pathway<br>Pathways in cancer<br>Plant-pathogen interaction<br>Progesterone-mediated oocyte maturation<br>Prostate cancer<br>Protein processing in endoplasmic reticulum |
| Creatine kinase B-type | Ckb | 62.7 | 42.713 | 311 | 0.36639595 | 2.572327037 | 1.3746 | ko00330 | Arginine and proline metabolism |
| Guanine nucleotide-binding protein G(I)/G(S)/G(T) subunit beta-2 | Gnb2 | 20.9 | 37.331 | 148 | -0.180434704 | -2.728495349 | 1.4652 | ko04062 | Chemokine signaling pathway |
| Synaptosomal-associated protein 25 | Snap25 | 71.4 | 23.315 | 276 | -0.303587437 | -2.584567189 | 1.3818 | ko04130 | SNARE interactions in vesicular transport |
| Poly(rC)-binding protein 1 | Pcbp1 | 12.1 | 37.497 | 68 | -0.339476585 | -5.161235546 | 2.6793 | ko03040 | Spliceosome |
| Sodium/potassium-transporting ATPase subunit alpha-1 | Atp1a1 | 25.8 | 112.98 | 395 | -0.356846809 | -2.626044796 | 1.4059 | ko04260<br>ko04960<br>ko04961<br>ko04964<br>ko04970<br>ko04971<br>ko04972<br>ko04973<br>ko04974<br>ko04976<br>ko04978 | Aldosterone-regulated sodium reabsorption<br>Bile secretion<br>Carbohydrate digestion and absorption<br>Cardiac muscle contraction<br>Endocrine and other factor-regulated calcium reabsorption<br>Gastric acid secretion<br>Mineral absorption<br>Pancreatic secretion<br>Protein digestion and absorption<br>Proximal tubule bicarbonate reclamation<br>Salivary secretion |
| Poly(rC)-binding protein 3 | Pcbp3 | 20.8 | 24.117 | 82 | -0.35725832 | -2.846988801 | 1.5331 | ko00250<br>ko04270<br>ko04510 | Alanine, aspartate and glutamate metabolism<br>Vascular smooth muscle contraction<br>Focal adhesion<br>Regulation of actin cytoskeleton |
|  |  |  |  |  |  |  |  | ko04810<br>ko04921 | Oxytocin signaling pathway |
| Guanine nucleotide-binding protein G(o) subunit alpha | Gnao1 | 37.6 | 40.084 | 201 | -0.388885021 | -2.47156149 | 1.3155 | ko04730<br>ko04916<br>ko05142<br>ko05145 | Chagas disease (American trypanosomiasis)<br>Long-term depression<br>Melanogenesis<br>Toxoplasmosis |
| Plasma membrane calcium-transporting ATPase 1 | Atp2b1 | 4.7 | 134.75 | 48 | -0.422261715 | -3.197607557 | 1.7291 | ko04020<br>ko04970<br>ko04972 | Calcium signaling pathway<br>Pancreatic secretion<br>Salivary secretion |
| ATP synthase subunit O, mitochondrial | Atp5o | 37.1 | 23.363 | 79 | -0.423535824 | -3.125266485 | 1.6893 | ko00190<br>ko05010<br>ko05012<br>ko05016 | Alzheimer's disease<br>Huntington's disease<br>Oxidative phosphorylation<br>Parkinson's disease |
| Guanine nucleotide-binding protein G(i) subunit alpha-2 | Gnai2 | 22.5 | 40.489 | 103 | -0.461163998 | -3.18320836 | 1.7212 | ko04020<br>ko04062<br>ko04360<br>ko04530<br>ko04540<br>ko04670<br>ko04730<br>ko04740<br>ko04914<br>ko04916<br>ko04971<br>ko05142<br>ko05145<br>ko05146 | Amoebiasis<br>Axon guidance<br>Calcium signaling pathway<br>Chagas disease (American trypanosomiasis)<br>Chemokine signaling pathway<br>Gap junction<br>Gastric acid secretion<br>Leukocyte transendothelial migration<br>Long-term depression<br>Melanogenesis<br>Olfactory transduction<br>Progesterone-mediated oocyte maturation<br>Tight junction<br>Toxoplasmosis |
| Vimentin | Vim | 73.6 | 53.687 | 584 | -0.532655716 | -2.526809502 | 1.3480 | ko05014<br>ko05410<br>ko05412<br>ko05414 | Amyotrophic lateral sclerosis (ALS)<br>Arrhythmogenic right ventricular cardiomyopathy (ARVC)<br>Dilated cardiomyopathy<br>Hypertrophic cardiomyopathy (HCM) |
| Mitochondrial glutamate carrier 1 | Slc25a22 | 26.6 | 24.64 | 27 | -0.689178944 | -3.51807772 | 1.9014 |  |  |
| Heterogeneous nuclear ribonucleoprotein A1 | Hnrnpa1 | 30.3 | 38.833 | 65 | -0.78055954 | -3.272488143 | 1.7700 | ko03040 | Spliceosome |
| Histone H1.2 | Hist1h1c | 21.2 | 21.266 | 115 | -1.250981331 | -2.995659744 | 1.6172 | ko05322<br>ko05034<br>ko04115<br>ko04914 | Systemic lupus erythematosus<br>Alcoholism<br>p53 signaling pathway<br>Progesterone-mediated oocyte maturation |
| T-complex protein 1 subunit theta | Cct8 | 10.8 | 53.082 | 21 | -1.38865757 | -3.498318032 | 1.8910 |  |  |
| Histone H1.4 | Hist1h1e | 16 | 21.977 | 119 | -1.393318176 | -3.013641476 | 1.6272 | ko05322<br>ko05034 | Systemic lupus erythematosus<br>Alcoholism |
| Histone H1.3 | Hist1h1d | 9.5 | 22.099 | 105 | -1.504905224 | -3.187679564 | 1.7237 | ko05322<br>ko05034 | Systemic lupus erythematosus<br>Alcoholism |
| Heterogeneous nuclear ribonucleoproteins C1/C2 | Hnrnpc | 28.8 | 34.384 | 51 | -1.816102505 | -5.20234665 | 2.6967 | ko03040 | Spliceosome |
| V-type proton ATPase subunit E 1 | Atp6v1e1 | 27.9 | 26.157 | 29 | -1.93034029 | -2.750957456 | 1.4782 | ko00190;<br>ko04145; | Collecting duct acid secretion<br>Epithelial cell signaling in Helicobacter pylori |
|  |  |  |  |  |  |  |  | ko04966;<br>ko05110;<br>ko05120;<br>ko05323 | infection<br>Oxidative phosphorylation<br>Phagosome<br>Rheumatoid arthritis<br>Vibrio cholerae infection |
| Heterogeneous<br>nuclear<br>ribonucleoprotein<br>U-like protein 2 | Hnrnpul2 | 16.1 | 84.939 | 40 | -1.97420311 | -3.755246915 | 2.0245 |  |  |
| Methyl-CpG-binding protein 2 | Mecp2 | 19 | 52.307 | 22 | -2.162194252 | -3.561647541 | 1.9243 | ko05016<br>ko04919<br>ko05202<br>ko04330<br>ko04110<br>ko04152 | Huntington s disease<br>Thyroid hormone signaling pathway<br>Transcriptional misregulation in cancer<br>Notch signaling pathway<br>Cell cycle<br>AMPK signaling pathway |
| Histone H1.5 | Hist1h1b | 15.7 | 22.576 | 26 | -2.227953911 | -3.3561123 | 1.8152 | ko05322<br>ko04115 | Systemic lupus erythematosus<br>p53 signaling pathway |
| Heterogeneous<br>nuclear<br>ribonucleoprotein<br>A/B | Hnrnpab | 29.6 | 33.816 | 30 | -2.522982121 | -3.297190543 | 1.7834 | ko03040 | Spliceosome |
| Sodium- and<br>chloride-dependent<br>GABA transporter 3 | Slc6a11 | 7.7 | 69.96 | 28 | -2.539692879 | -3.292265714 | 1.7807 |  |  |
| Heterogeneous<br>nuclear<br>ribonucleoprotein<br>D0 | Hnrnpd | 22.7 | 24.611 | 50 | -2.972328186 | -3.282508326 | 1.7754 | ko03040 | Spliceosome |
| Histone H1.1 | Hist1h1a | 23 | 21.785 | 26 | -3.279434204 | -4.172174855 | 2.2317 | 04115<br>04110<br>05322<br>05214<br>05200<br>04068 | p53 signaling pathway<br>Cell cycle<br>Systemic lupus erythematosus<br>Glioma<br>Pathways in cancer<br>FoxO signaling pathway |

The Akita vs C57 combination up and down regulated proteins were involved in various molecular functions, biological process, cellular components, protein classes and biological pathways. The up and down regulated proteins were involved in various biological processes such as biological regulation (GO:0065007), cellular component organization or biogenesis (GO:0071840), cellular process (GO:0009987), developmental process (GO:0032502), localization (GO:0051179), metabolic process (GO:0008152), multicellular organismal process (GO:0032501), response to stimulus (GO:0050896) respectively. Nme1, Prdx6, Gm20390, Nme2, Atp1a1, Atp1b2, Atp1a2 proteins are involved in biological regulation (GO:0065007); Epb4.1l3, Lmnb2, Map2, Cfl1 proteins are involved in cellular component organization or biogenesis (GO:0071840); Hnrnpk, Pcbp1, Gnb1, Epb4.1l3, Nme1, Uqcrq, Map2, Prdx6, Gnb2, Gm20390, Atp6v1a, Nme2, Cfl1, Atp1a1, Atp1b2, Atp1a2 proteins are involved in cellular process (GO:0009987); Hnrnpk, Pcbp1, Nme1, Map2, Gm20390, Nme2 proteins are involved in developmental process (GO:0032502); Hnrnpk, Pcbp1, Stxbp1 proteins are involved in localization (GO:0051179); Hnrnpk, Pcbp1, Nme1, Uqcrq, Gm20390, Prdx6, Atp6v1a, Nme2, Atp1a1, Glul, Atp1b2, Atp1a2 proteins are involved in metabolic process (GO:0008152); Hnrnpk, Pcbp1, Gnb1, Map2, Gnb2, Stxbp1 proteins are involved in multicellular organismal process (GO:0032501); Prdx6, Prdx5 proteins are involved in response to stimulus (GO:0050896) respectively (**Figure 4B**). The up and down regulated proteins are involved in various molecular functions such as antioxidant activity (GO:0016209) (Prdx6 and Prdx5), binding (GO:0005488) (Hnrnpk, Pcbp1, Gnb1, Map2, Gnb2 and Cfl1), catalytic activity (GO:0003824) (Hnrnpk, Pcbp1, Gnb1, Nme1, Uqcrq, Gm20390, Prdx6, Atp6v1a, Nme2, Atp1a1, Glul, Atp1b2, Atp1a2), structural molecule activity (GO:0005198) (Epb4.1l3 and Cfl1) and transporter activity (GO:0005215) (Uqcrq, Atp6v1a, Stxbp1, Atp1a1, Atp1b2, Atp1a2) respectively (**Figure 4C**). The DEPs are involved in various cellular component processes such as cell part (GO:0044464) (Epb4.1l3, Uqcrq, Map2, Prdx6, Atp6v1a, Cfl1), macromolecular complex (GO:0032991) (Hnrnpk, Pcbp1, Gnb1, Uqcrq, Gnb2 and Atp1b2), membrane (GO:0016020) (Uqcrq, Atp6v1a, Atp1ab2), andorganelle (GO: 0043226) (Epb4.1l3, Atp6v1a, Cfl1) respectively (**Figure 4D**). The DEPs belong to various protein classes such as cytoskeletal protein (Cfl1), enzyme modulator (Hnrnpk, Pcbp1, Gnb1 and Gnb2), hydrolase (Hnrnpk, Pcbp1, Gnb1 and Gnb2, Atp6v1a, Atp1a1 and Atp1a2), ligase (Glul), membrane traffic protein (Stxbp1), nucleic acid binding (Hnrnpk, Pcbp1 and Atp6v1a), oxidoreductase (Prdx6), transporter (Atp6v1a, Stxbp1, Atp1a1, Atp1b2 and Atp1a2) respectively (**Figure 4E**). We have also analyzed the reactome pathways of up and down regulated proteins, which are Ion transport by P-type ATPases (Atp1a1, Atp1b2 and Atp1a2), Ion channel transport (Atp6v1a, Atp1a1, Atp1b2 and Atp1a2), Transmembrane transport of small molecules (Gnb1, Gnb2, Atp6v1a, Atp1a1, Atp1b2 and Atp1a2), Ion homeostasis (Atp1a1, Atp1b2 and Atp1a2), Cardiac conduction (Atp1a1, Atp1b2 and Atp1a2), Regulation of insulin secretion (Gnb1, Gnb2 and Stxbp1), Integration of energy metabolism (Gnb1, Gnb2 and Stxbp1), mRNA Splicing - Major Pathway (Hnrnpk, Hnrnpa1, Pcbp1, Hnrnpc, and Hnrnpd), Processing of Capped Intron-Containing Pre-mRNA (Hnrnpk, Hnrnpa1, Pcbp1, Hnrnpc, and Hnrnpd), Transmission across Chemical Synapses (Gnb1, Gnb2, Stxbp1 and Glul), Neuronal System (Gnb1, Epb4.1l3, Gnb2, Stxbp1 and Glul) and Unclassified pathways (Lmnb2, Hnrnpab, Nefm, Gm20390, Map2, Ntm and Nme2) respectively (**Figure 4F**). **Table 3** showed the up and down regulated proteins, gene name, MS/MS count, unique sequence coverage, molecular weight (MW), T-test difference, p-value, various KEGG name and pathways.

**Table 3:**
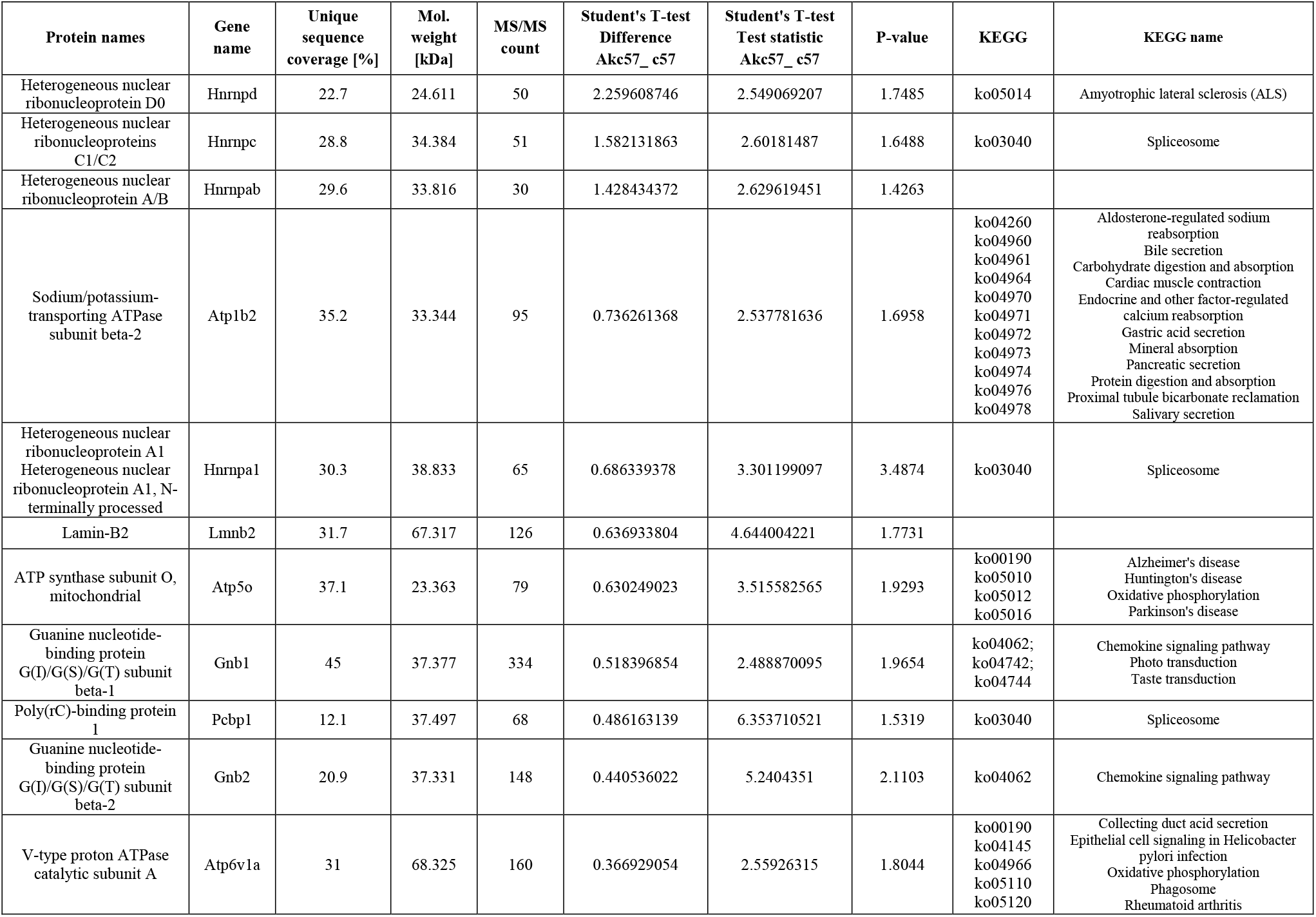
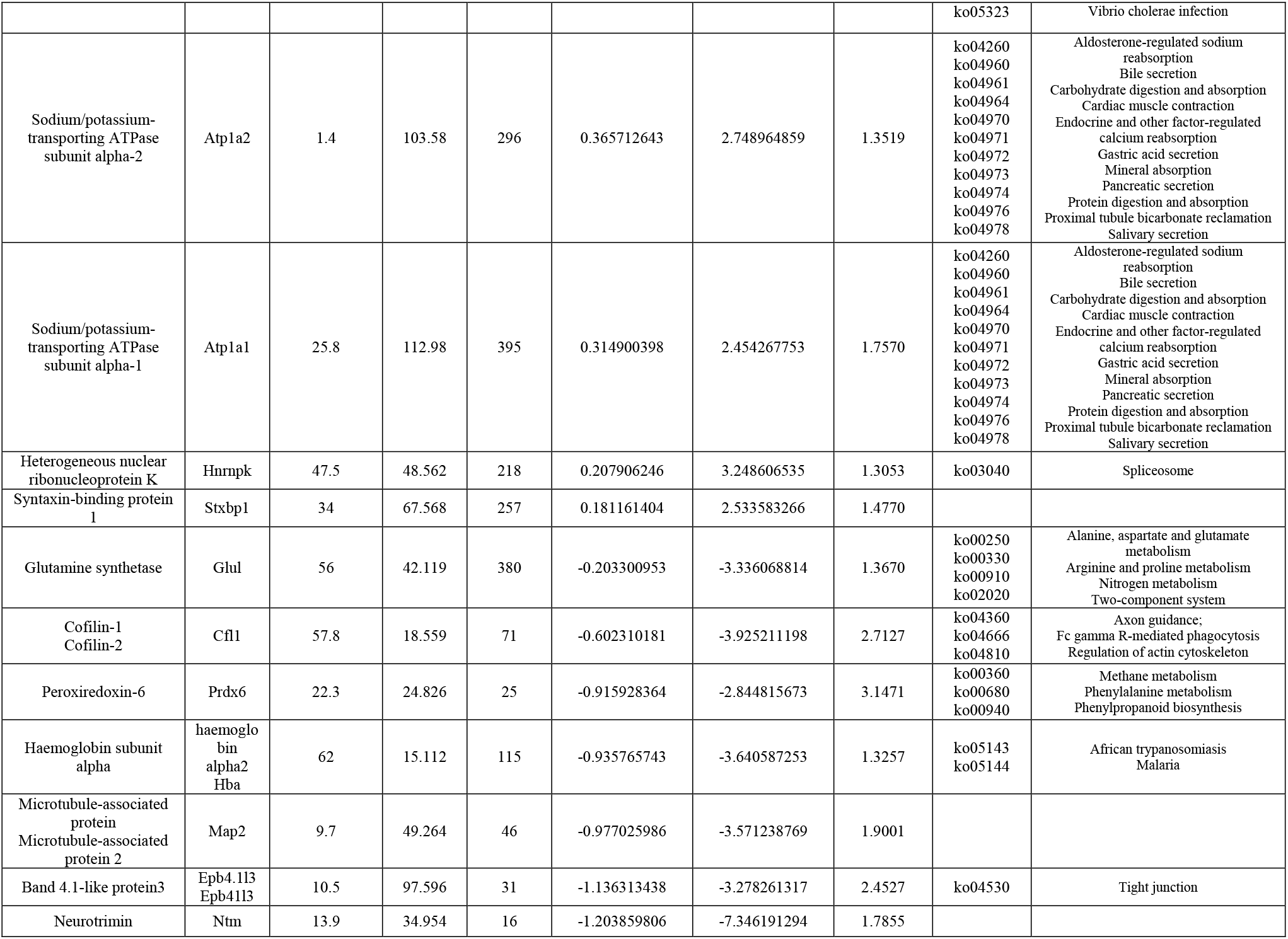
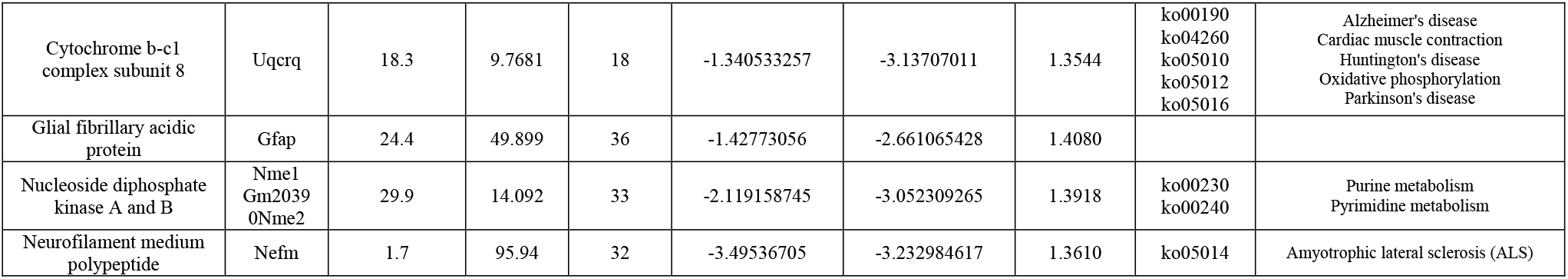
The list of significantly differentially expressed proteins, gene name, MS/MS count, unique sequence coverage, molecular weight (MW), T-test difference, p-value, various KEGG name, and pathways in Akita group compared to C57 group.

The P1GF^−/−^ vs. C57 combination up and down-regulated proteins were involved in various molecular functions, biological process, cellular components, protein classes and biological pathways. The DEPs are involved various biological processes such as biological regulation (GO:0065007), cellular component organization or biogenesis (GO:0071840), cellular process (GO:0009987), developmental process (GO:0032502), localization (GO:0051179), locomotion (GO:0040011), metabolic process (GO:0008152), multicellular organismal process (GO:0032501), response to stimulus (GO:0050896) respectively. Napb, Rac3, Gnat1, Gnat3, Rhobtb3, Rac2, Rac1, Gnat2, Atp1b2 proteins are involved in biological regulation (GO:0065007); Napb, Rac3, Tubb3, Rhobtb3, Epb4.1l3, Rac1, Lmnb2, Tpm1, Tpm2 proteins are involved in cellular component organization or biogenesis (GO:0071840); Napb, Rac3, Gnat1, Gnb1, Gnat3, Tubb3, Ywhab, Rhobtb3, Epb4.1l3, Rac2, Gnb4, Gngt1, Tpm1, Rac1, Gnb2, Rac2, Gnat2, Tpm2, Nono, Crocc, Atp1b2, Aco2, Ywhag are involved in cellular process (GO:0009987); Rac1 and Rac2 are involved in developmental process (GO:0032502); Napb and Rac3 are involved in localization (GO:0051179); Rac3 is also involved in locomotion (GO:0040011); Srsf9, Rac3, Nono, Rhobtb3, Rac3, Gnat3, Aco2, Atp1b2, Cirbp, Gnat1, Gnat2, Rac1, Rac2, Srsf1 are involved in metabolic process (GO:0008152); Gnat3, Gnb1, Gnb2, Gnb4, Rac1, Rac2, Napb, Tpm1, Tpm2 are involved in multicellular organismal process (GO:0032501), and Gnat1, Gnat2, Gnat3, Rac3, Rac2, Rac1, Rhobtb3 are involved in response to stimulus (GO:0050896) respectively (**Figure 5B**). The DEPs are involved in various molecular functions such as binding (GO:0005488) (Aco2, Cirbp, Crocc, Gnat1, Gnat2, Gnat3, Gnb1, Gnb2, Gnb4, Gngt1, Napb, Nono, Rac1, Rac2, Rac3, Rhobtb3, Srsf1, Srsf9, Tmp1, Tmp2 and Tubb3), catalytic activity (GO:0003824) (Aco2, Cirbp, Crocc, Gnat1, Gnat2, Gnat3, Gnb1, Gnb2, Gnb4, Gngt1, Napb, Nono, Rac1, Rac2, Rac3, Rhobtb3, Srsf1, Srsf9), signal transducer activity (GO:0004871) (Gnat1, Gnat2, Gnat3, Rac1, Rac2, Rac3), structural molecule activity (GO:0005198) (Epb4.1l3 and Tubb3), transporter activity (GO:0005215) (Atp1b2) respectively (**Figure 5C**). The DEPs are involved in cellular component process such as cell part (GO:0044464) (Aco2, Epb4.1l3, Gnat1, Gnat2, Gnat3, Napb, Rac3, Ndufa8, Nono, Rhobtb3, Tmp1, Tmp2, and Tubb3), macromolecular complex (GO:0032991) (Atp1b2, Gnat1, Gnat2, Gnat3, Gnb1, Gnb2, Gnb4, Gngt1, Napb, Ndufa8, Tpm1, and Tpm2), membrane (GO:0016020) (Atp1b2, Gnat1, Gnat2, Gnat3, Napb, Ndufa8, Rac3 and Rhobtb3), organelle (GO:0043226) (Aco2, Epb4.1l3, Napb, Rac3, Nono, Tpm1, Tpm2, Rhobtb3 and Tubb3), and synapse (GO:0045202) (Napb) respectively (**Figure 5D**). The DEPs are classified into various protein class such as chaperone (Ywhab and Ywhag), cytoskeletal protein (Tubb3, Tpm1 and Tpm2), enzyme modulator (Rac3, Gnat1, Gnb1, Gnat3, Rhobtb3, Rac2, Gnb4, Gngt1, Rac1, Gnb2, Gnat2 and Crocc), hydrolase (Gnb1, Gnb2, Gnb4), lyase (Aco2), membrane traffic protein (Napb), nucleic acid binding (Cirbp, Srsf9, Srsf1 and Nono), oxidoreductase (Ndufa8), transcription factor (Rac3), transferase (Rac3), transporter (Atp1b2), and viral protein (Crocc) respectively (**Figure 5E**). We have also analyzed the reactome pathways of up and down regulated proteins, which are Activation of the phototransduction cascade (Gnat1, Gnb1 and Gnat2), Signal Transduction (Rac3, Gnat1, Gnb1, Gnat3, Tubb3, Ywhab, Rac2, Gnb4, Gngt1, Rac1, Gnb2, Gnat2, Gfap and Ywhag), G-protein activation (Gnat1, Gnb1, Gnat3 and Gnat2), Opioid Signalling (Gnat1, Gnb1, Gnat3 and Gnat2), Signalling by GPCR (Gnat1, Gnb1, Gnat3, Ywhab, Rac2, Gnb4, Gngt1, Gnb2, Rac1 and Gnat2), ADP signalling through P2Y purinoceptor-12 (Gnb1 and Gnat3), Platelet activation, signalling and aggregation (Gnb1, Gnat3, Rac2 and Rac1), Hemostasis (Gnb1, Gnat3, Rac2 and Atp1b2), Cooperation of PDCL (PhLP1) and TRiC/CCT in G-protein beta folding (Gnat1, Gnb1, Gnat3, Gnb4, Gnb2 and Gnat2), Chaperonin-mediated protein folding (Gnat1, Gnb1, Gnat3, Gnb4, Gnb2 and Gnat2), Metabolism of proteins (Gnat1, Gnb1, Gnat3, Eif4a1, Gnb4, Eif4a2, Gnb2, Hnrnpc and Gnat2), Activation of BAD and translocation to mitochondria (Ywhab and Ywhag), Intrinsic Pathway for Apoptosis (Ywhab and Ywhag) Adrenaline, noradrenaline inhibits insulin secretion (Gnb1, Gnb2, Gnb4 and Gngt1), Regulation of insulin secretion (Gnb1, Vamp2, Gnb2, Gnb4 and Gngt1), Metabolism (Atp5o, Gnb1, Vamp2, Ndufa8, Atp5f1, Gnb4, Gngt1, Gnb2, Aco2 and Hba), RHO GTPase Effectors (Ywhab, Rac2, Rac1 and Ywhag), Signaling by Rho GTPases (Ywhab, Rac2, Rac1 and Ywhag), Glucagon-type ligand receptors (Gnb1, Gnb2, Gnb4 and Gngt1), Activation of G protein gated Potassium channels (Gnb1, Gnb2, and Gngt1), Transmission across Chemical Synapses (Gnb1, Gnat3, Vamp2, Gngt1 and Gnb2), Glucagon-like Peptide-1 (GLP1) regulates insulin secretion (Gnb1, Gnb2, Gnb4 and Gngt1), Inhibition of voltage gated Ca2+ channels via Gbeta/gamma subunits (Gnb1, Gnb2, and Gngt1), Neuronal System (Gnb1, Gnat3, Vamp2, Epb4.1l3, Gngt1 and Gnb2), Inactivation, recovery and regulation of the phototransduction cascade (Gnat1 and Gngt1), Beta-catenin independent WNT signalling (Gnb1, Rac1 and Gnat2), Rho GTPase cycle (Rac1, Rac2, and Rac3) and un classified pathway (Cirbp, Napb, Vamp1, Lmnb2, Nefm, Nono, Crocc and Rbp3) respectively (**Figure 5F**). **Table 4** showed the up and down-regulated proteins, gene name, MS/MS count, unique sequence coverage, molecular weight (MW), T-test difference, p-value, various KEGG name, and pathways.

**Table 4:**
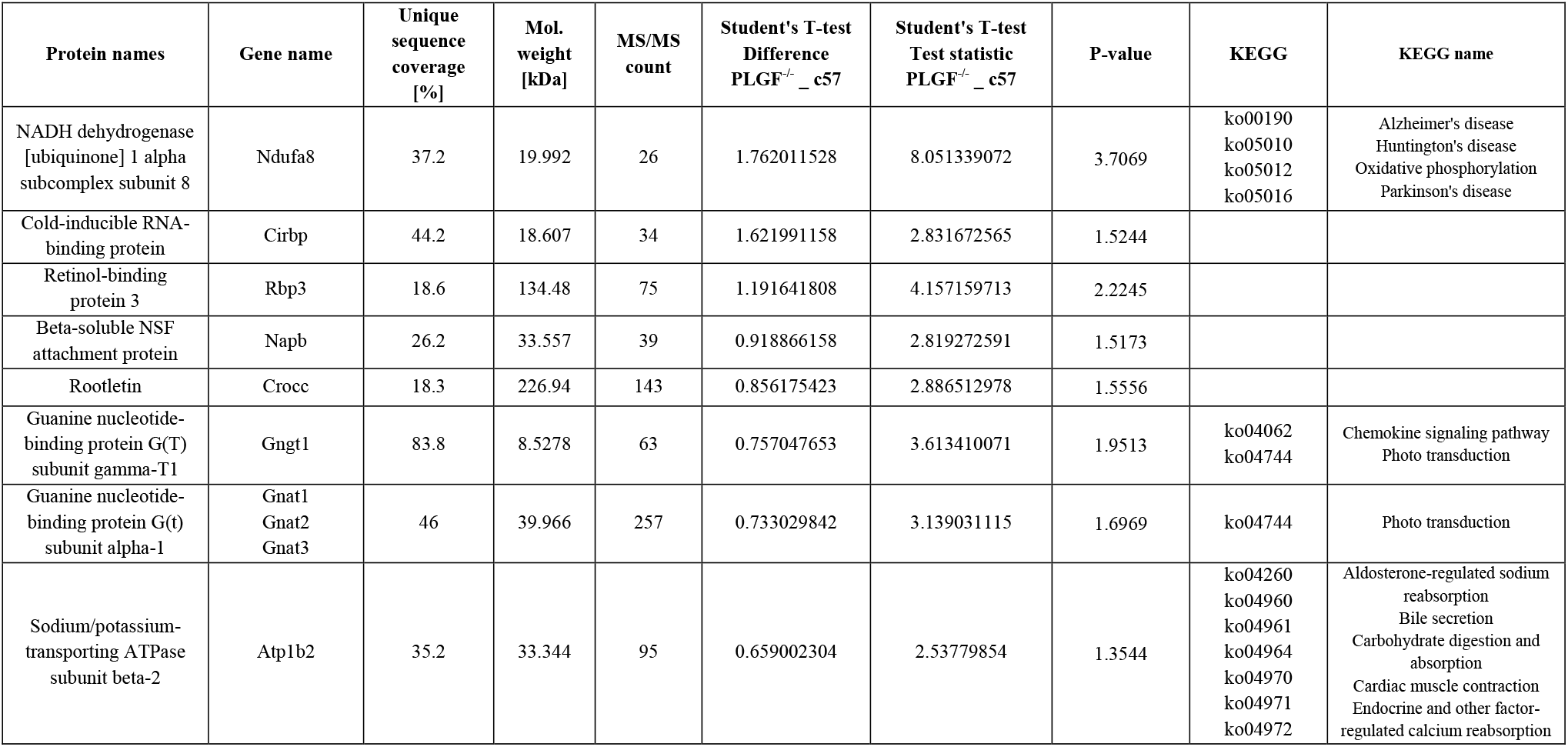
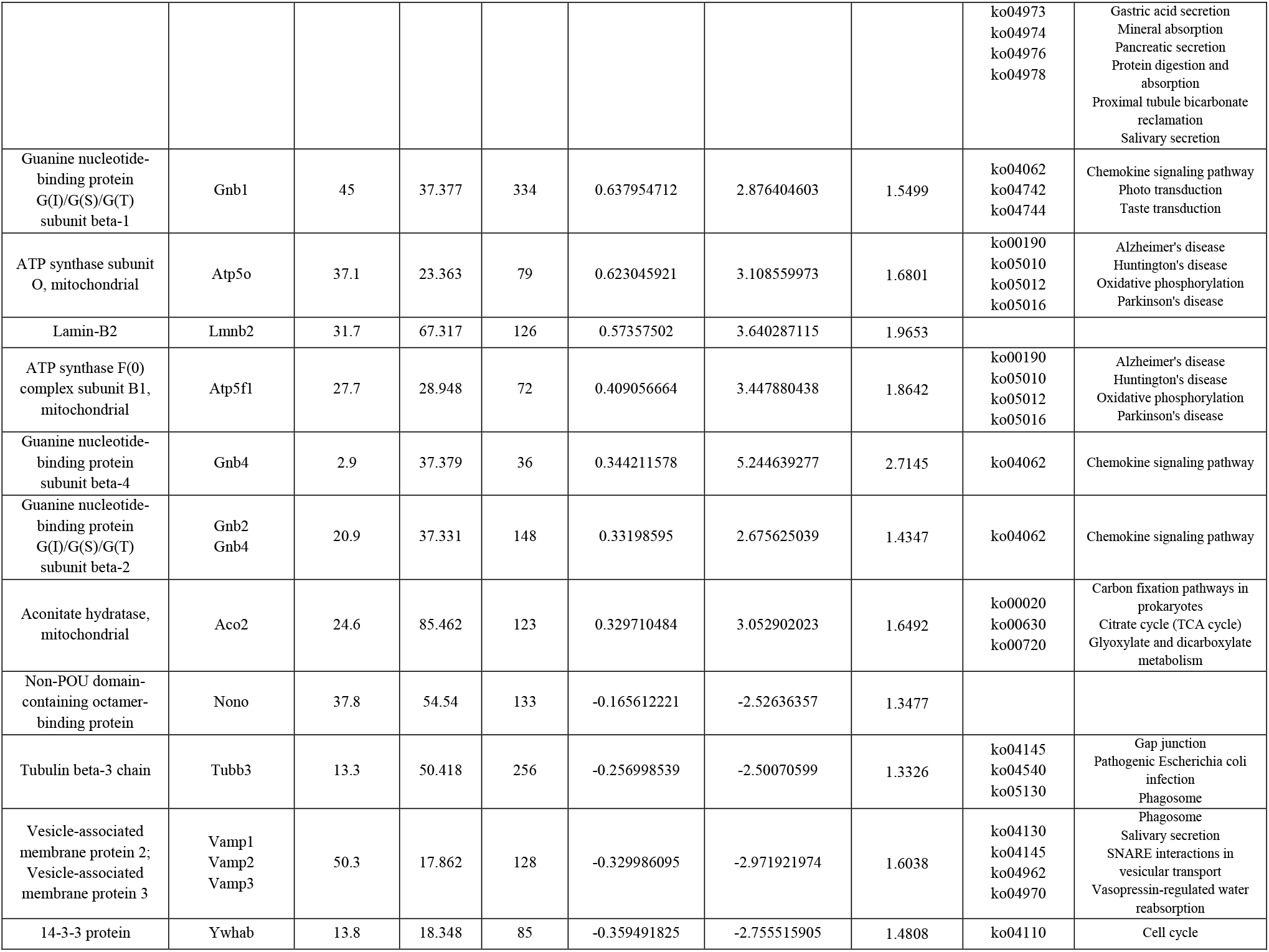
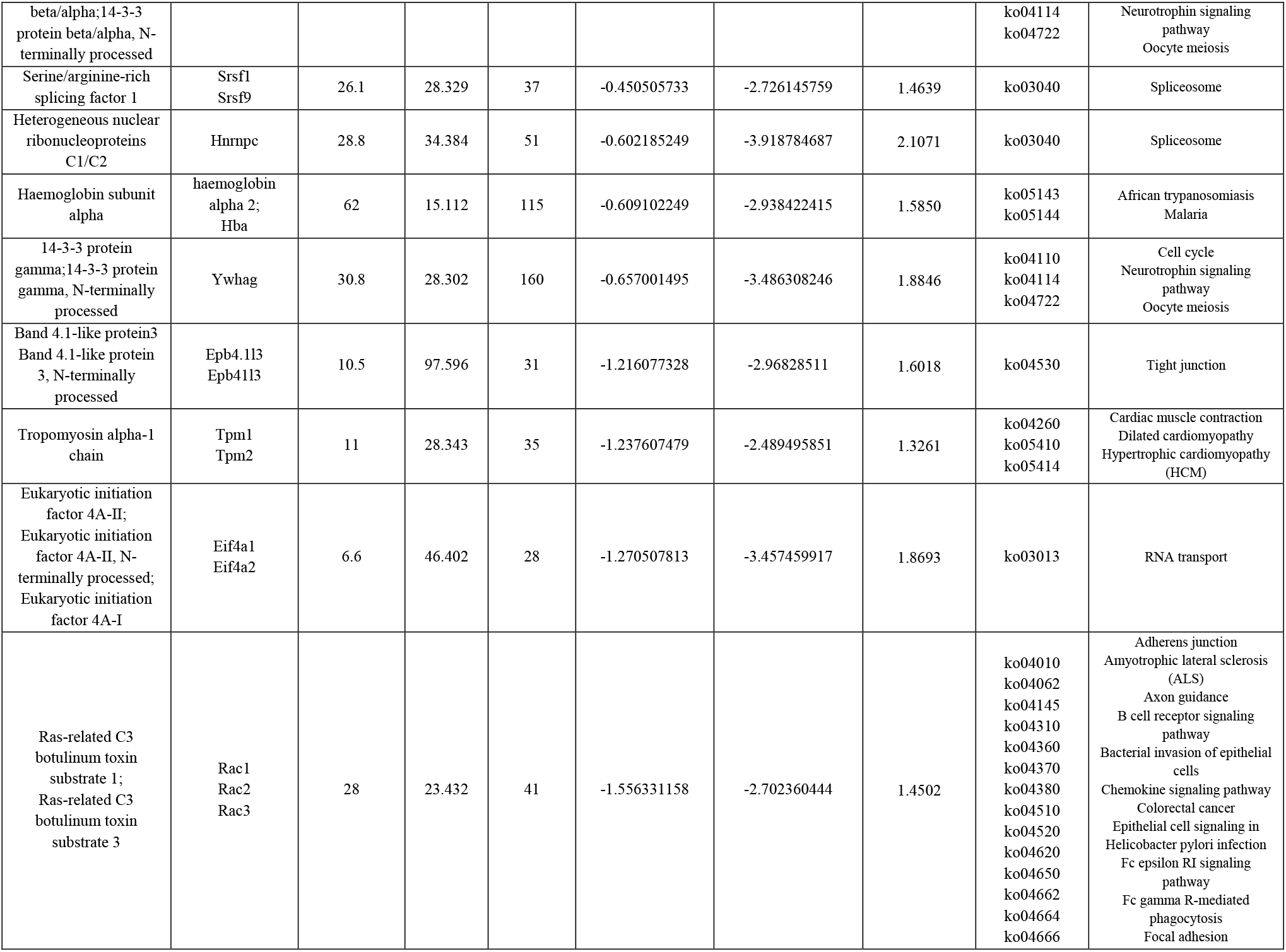
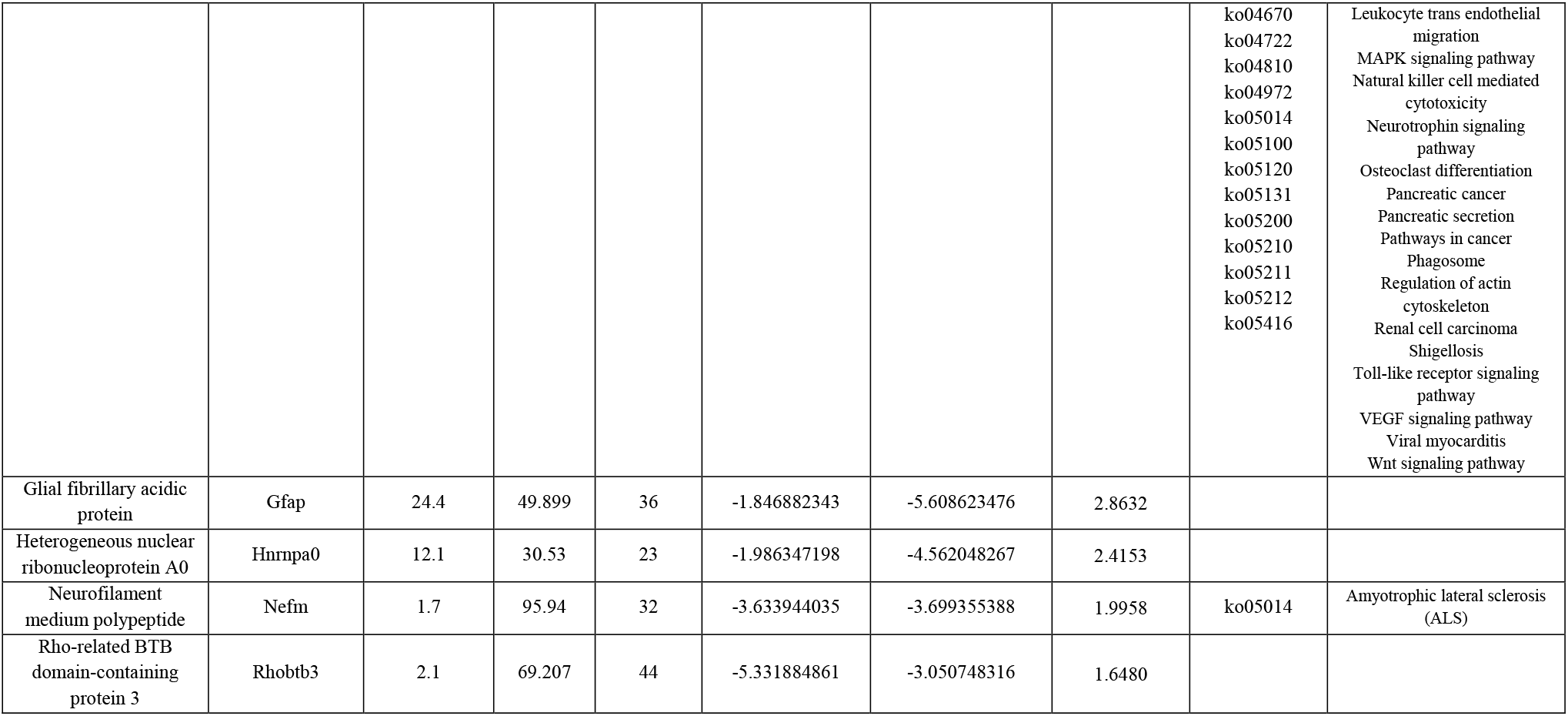
The list of significantly differentially expressed proteins, gene name, MS/MS count, unique sequence coverage, molecular weight (MW), T-test difference, p-value, various KEGG name, and pathways in PLGF^−/−^ group compared to C57 group.

The Akita.P1GF^−/−^ vs. P1GF^−/−^ combination up and down-regulated proteins are involved in various molecular functions, biological processes, cellular components, protein classes and biological pathways. The DEPs are involved in various biological process such as biological regulation (GO:0065007), cellular component organization or biogenesis (GO:0071840), cellular process (GO:0009987), developmental process (GO:0032502), localization (GO:0051179), metabolic process (GO:0008152), multicellular organismal process (GO:0032501), and response to stimulus (GO:0050896) respectively. Map1b and Hnrnpm are involved in biological regulation (GO:0065007), Tubb2b, Tubb3, Tubb6, Cct8, Map1b, and Tubb5 are involved in cellular component organization or biogenesis (GO:0071840), Hspa8, Tubb2b, Tubb3, Tubb6, Cox5b, Cct8, Map1b, Hnrnpm, Tubb5, and Pura proteins are involved in cellar process (GO:0009987), Map1b is involved in developmental process (GO:0032502), Hspa8, Vdac2 and Cox5b are involved in localization (GO:0051179) biological process, Cirbp, Hspa8, Hsp90ab1, Cox5b, Cct8, Erh, Hnrnpm and Pura protein are involved in metabolic process (GO:0008152), Map1b is involved in multicellular organismal process (GO:0032501), and Hspa8, Hsp90ab1, Mecp2, Hnrnpm are involved in response to stimulus (GO:0050896) respectively (**Figure 6B**). The DEPs were involved in various molecular functions such as binding (GO:0005488) (Cirbp, Hspa8, Tubb2b, Tubb3, Mecp2, Tubb6, Cct8, Map1b, Erh, Hnrnpm, Tubb5 and Pura), catalytic activity (GO:0003824) (Cirbp, Hspa8, and Cox5b), structural molecule activity (GO:0005198) (Tubb2b, Tubb3, Tubb6, and Tubb5), and transporter activity (GO:0005215) (Vdac2 and Cox5b) respectively (**Figure 6C**). The DEPs were also involved in various cellular component functions such as (**Figure 6D**). The DEPs are belonged to various protein classes such as cell adhesion molecule (Rs1), chaperone (Hsp90ab1, Cct8), cytoskeletal protein (Tubb2b, Tubb3, Tubb6, Map1b and Tubb5), hydrolase (Rs1), nucleic acid binding (Cirbp, Mecp2, Hnrnpm, Pura), oxidoreductase (Ndufa8, Cox5b), receptor (Rs1), signalling molecule (Rs1), transcription factor (Erh, Pura), transferase (Mecp2) and transporter (Vdac2) respectively (**Figure 6E**). The DEPs are also involved in various reactome pathways such as Attenuation phase (Hspa8 and Hsp90ab1), HSF1-dependent transactivation (Hspa8 and Hsp90ab1), Intraflagellar transport (Tubb2b, Tubb3, and Tubb6), Assembly of the primary cilium (Tubb2b, Tubb3, Tubb6, and Tubb5), Organelle biogenesis and maintenance (Tubb2b, Tubb3, Tubb6, and Tubb5), Translocation of GLUT4 to the plasma membrane (Tubb2b, Tubb3, and Tubb6), Hedgehog ‘off’ state (Tubb2b, Tubb3, and Tubb6), and Hedgehog ‘on’ state (Tubb2b, Tubb3, and Tubb6) respectively (**Figure 6F**). **Table 5** showed the up and down-regulated proteins, gene name, MS/MS count, unique sequence coverage, molecular weight (MW), T-test difference, p-value, various KEGG name, and pathways.

**Table 5:**
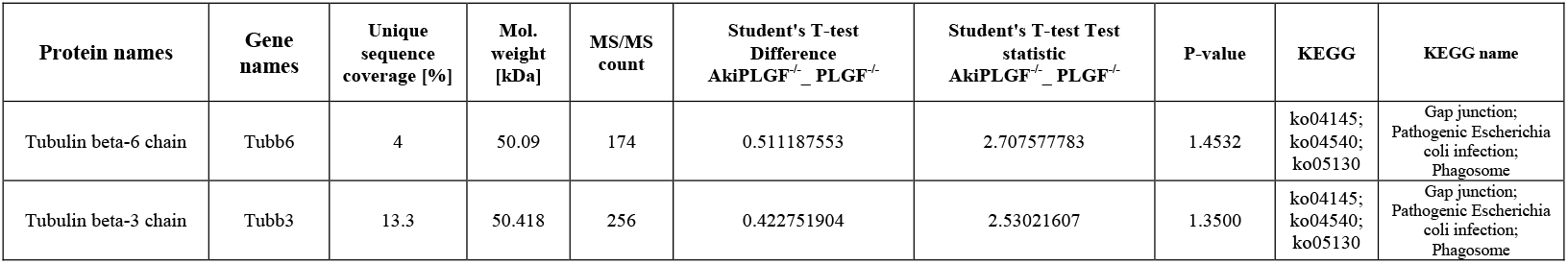
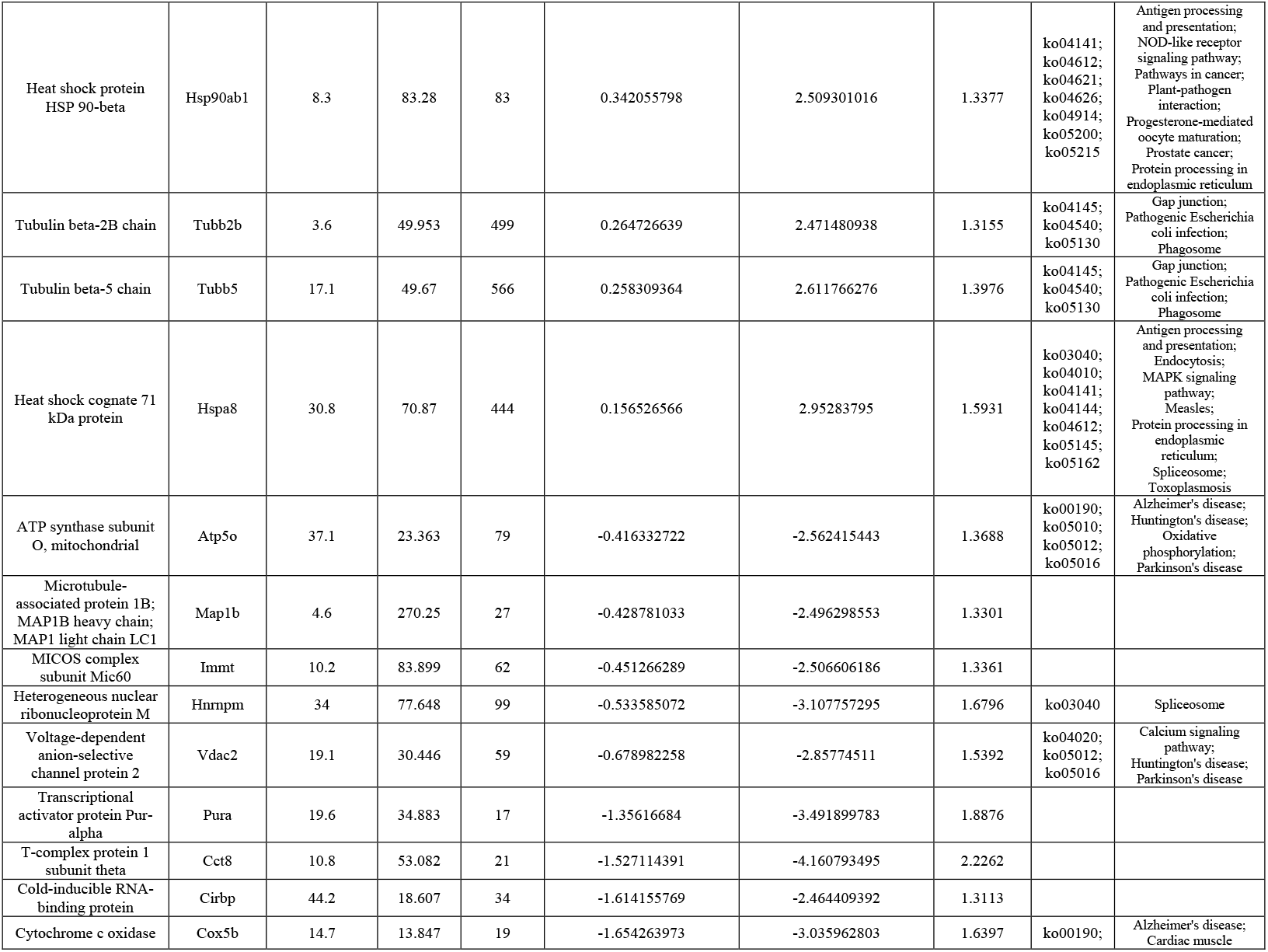
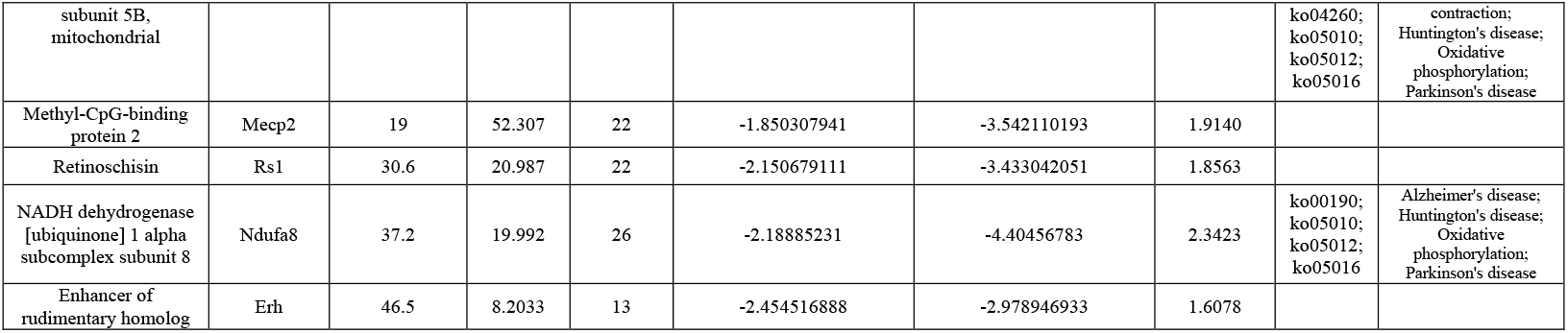
The list of significantly differentially expressed proteins, gene name, MS/MS count, unique sequence coverage, molecular weight (MW), T-test difference, p-value, various KEGG name, and pathways in Akita.P1GF^−/−^ group compared to P1GF^−/−^ group.

| Protein names | Gene names | Unique sequence coverage [%] | Mol. weight [kDa] | MS/MS count | Student's T-test Difference AkiPLGF<sup>-/-</sup>_ PLGF<sup>-/-</sup> | Student's T-test Test statistic AkiPLGF<sup>-/-</sup>_ PLGF<sup>-/-</sup> | P-value | KEGG | KEGG name |
| --- | --- | --- | --- | --- | --- | --- | --- | --- | --- |
| Tubulin beta-6 chain | Tubb6 | 4 | 50.09 | 174 | 0.511187553 | 2.707577783 | 1.4532 | ko04145;<br>ko04540;<br>ko05130 | Gap junction;<br>Pathogenic Escherichia coli infection;<br>Phagosome |
| Tubulin beta-3 chain | Tubb3 | 13.3 | 50.418 | 256 | 0.422751904 | 2.53021607 | 1.3500 | ko04145;<br>ko04540;<br>ko05130 | Gap junction;<br>Pathogenic Escherichia coli infection;<br>Phagosome |
| Heat shock protein HSP 90-beta | Hsp90ab1 | 8.3 | 83.28 | 83 | 0.342055798 | 2.509301016 | 1.3377 | ko04141;<br>ko04612;<br>ko04621;<br>ko04626;<br>ko04914;<br>ko05200;<br>ko05215 | Antigen processing and presentation;<br>NOD-like receptor signaling pathway;<br>Pathways in cancer;<br>Plant-pathogen interaction;<br>Progesterone-mediated oocyte maturation;<br>Prostate cancer;<br>Protein processing in endoplasmic reticulum |
| Tubulin beta-2B chain | Tubb2b | 3.6 | 49.953 | 499 | 0.264726639 | 2.471480938 | 1.3155 | ko04145;<br>ko04540;<br>ko05130 | Gap junction;<br>Pathogenic Escherichia coli infection;<br>Phagosome |
| Tubulin beta-5 chain | Tubb5 | 17.1 | 49.67 | 566 | 0.258309364 | 2.611766276 | 1.3976 | ko04145;<br>ko04540;<br>ko05130 | Gap junction;<br>Pathogenic Escherichia coli infection;<br>Phagosome |
| Heat shock cognate 71 kDa protein | Hspa8 | 30.8 | 70.87 | 444 | 0.156526566 | 2.95283795 | 1.5931 | ko03040;<br>ko04010;<br>ko04141;<br>ko04144;<br>ko04612;<br>ko05145;<br>ko05162 | Antigen processing and presentation;<br>Endocytosis;<br>MAPK signaling pathway;<br>Measles;<br>Protein processing in endoplasmic reticulum;<br>Spliceosome;<br>Toxoplasmosis |
| ATP synthase subunit O, mitochondrial | Atp5o | 37.1 | 23.363 | 79 | -0.416332722 | -2.562415443 | 1.3688 | ko00190;<br>ko05010;<br>ko05012;<br>ko05016 | Alzheimer's disease;<br>Huntington's disease;<br>Oxidative phosphorylation;<br>Parkinson's disease |
| Microtubule-associated protein 1B; MAP1B heavy chain; MAP1 light chain LC1 | Map1b | 4.6 | 270.25 | 27 | -0.428781033 | -2.496298553 | 1.3301 |  |  |
| MICOS complex subunit Mic60 | Immt | 10.2 | 83.899 | 62 | -0.451266289 | -2.506606186 | 1.3361 |  |  |
| Heterogeneous nuclear ribonucleoprotein M | Hnrnpm | 34 | 77.648 | 99 | -0.533585072 | -3.107757295 | 1.6796 | ko03040 | Spliceosome |
| Voltage-dependent anion-selective channel protein 2 | Vdac2 | 19.1 | 30.446 | 59 | -0.678982258 | -2.85774511 | 1.5392 | ko04020;<br>ko05012;<br>ko05016 | Calcium signaling pathway;<br>Huntington's disease;<br>Parkinson's disease |
| Transcriptional activator protein Pur-alpha | Pura | 19.6 | 34.883 | 17 | -1.35616684 | -3.491899783 | 1.8876 |  |  |
| T-complex protein 1 subunit theta | Cct8 | 10.8 | 53.082 | 21 | -1.527114391 | -4.160793495 | 2.2262 |  |  |
| Cold-inducible RNA-binding protein | Cirbp | 44.2 | 18.607 | 34 | -1.614155769 | -2.464409392 | 1.3113 |  |  |
| Cytochrome c oxidase | Cox5b | 14.7 | 13.847 | 19 | -1.654263973 | -3.035962803 | 1.6397 | ko00190; | Alzheimer's disease;<br>Cardiac muscle |
| subunit 5B,<br>mitochondrial |  |  |  |  |  |  |  | ko04260;<br>ko05010;<br>ko05012;<br>ko05016 | contraction;<br>Huntington's disease;<br>Oxidative<br>phosphorylation;<br>Parkinson's disease |
| Methyl-CpG-binding<br>protein 2 | Mecp2 | 19 | 52.307 | 22 | -1.850307941 | -3.542110193 | 1.9140 |  |  |
| Retinoschisin | Rs1 | 30.6 | 20.987 | 22 | -2.150679111 | -3.433042051 | 1.8563 |  |  |
| NADH dehydrogenase<br>[ubiquinone] 1 alpha<br>subcomplex subunit 8 | Ndufa8 | 37.2 | 19.992 | 26 | -2.18885231 | -4.40456783 | 2.3423 | ko00190;<br>ko05010;<br>ko05012;<br>ko05016 | Alzheimer's disease;<br>Huntington's disease;<br>Oxidative<br>phosphorylation;<br>Parkinson's disease |
| Enhancer of<br>rudimentary homolog | Erh | 46.5 | 8.2033 | 13 | -2.454516888 | -2.978946933 | 1.6078 |  |  |

#### Protein-protein network analysis

The protein-protein network analysis is a wide-ranging approach to know the annotation of desire proteome (Kumari et al., 2015). The functional network protein study will be helpful for drug discovery, to understand metabolic pathways and to predict or develop genotype-phenotype associations (Wang and Moult, 2001; Wang et al., 2009). We have performed the protein-protein network analysis for all DEPs using STRING 10 database (https://string-db.org/). The DEPs of Akita.P1GF^−/−^ vs. Akita have uploaded to STRING network set the mouse database. They have total 30 nodes, 26 edges, cluster coefficient 0.497, average node degree:1.73, PPI enrichment p-value:1.17e-05. In KEGG pathway network, Gnb2, Gnai2, and Gnao1 are involved in Retrograde endocannabinoid signaling pathway (ko04723), Glutamatergic synapse pathway (ko04724), Cholinergic synapse pathway (ko04725), GABAergic synapse pathway (ko04727), Dopaminergic synapse pathway (ko04728), Serotonergic synapse pathway (ko04726). Pcb1, Hnrnpc, and Hnrnpa1 are involved in Spliceosome pathway (ko03040), Snap25 and Atp6v1e1 are involved in Synaptic vesicle cycle pathway (ko04721). **Figure 3G** highlighted the up and down-regulated proteins are involved in complex protein assembly (yellow) in biological process (GO:0006461), binding (blue) function of molecular function (GO:0005488), membrane-bounded organelle (red) proteins in cellular component (GO:0043227). The Akita vs. C57 up, and down-regulated proteins have total 28 nodes, 27 edges, cluster co-efficient 0.539, average node degree:1.73, PPI enrichment p-value: 2.62e-05. **Figure 4G** highlighted the up and down-regulated proteins involved in nervous system development (GO:0007399) (red), response to glucose (GO:0009749) (blue) of biological process, catalytic activity (GO:0003824) (light green), hydrolase activity (GO:0016787) (pink), of molecular functions, Insulin secretion (ko04911) (yellow), Pancreatic secretion (ko04972) (cyan), Oxidative phosphorylation (ko00190) (thick green) of KEGG pathways. The P1GF^−/−^ vs. C57 up, and down-regulated proteins have total 40 nodes, 87 edges, cluster co-efficient 0.692, average node degree:4.35, PPI enrichment p-value: 9.93e-10. **Figure 5G** highlighted the up and down regulated proteins involved in nervous system development (GO:0007399) (red), eye photoreceptor cell differentiation (GO:0001754) (blue) of biological process, hydrolase activity (GO:0016787) (light green), catalytic activity (GO:0003824) (yellow) of molecular functions, photoreceptor inner segment (GO:0001917) (pink), photoreceptor outer segment (GO:0001750) (thick green) of cellular components, Ras signalling pathway (ko04014) (cyan), VEGF signalling pathway (ko04370) (orange), Oxidative phosphorylation (ko00190) (magenta) of KEGG pathways. The Akita.P1GF^−/−^ vs P1GF^−/−^ up and down-regulated proteins have total 19 nodes, 14 edges, cluster co-efficient 0.511, average node degree:1.47, PPI enrichment p-value: 0.0013. **Figure 6G** highlighted the up and down-regulated proteins involved in protein folding (GO:0006457) (blue) of biological process, binding (GO:0005488) (red) of molecular function, intracellular organelle part (GO:0044446) (green) of cellular component, Gap junction (ko04540) (green) of KEGG pathway. The three and four combinations of up and down-regulated proteins are represented as a Venn diagram (**Figure 7A, B**), Akita vs. C57 have 30 up, and down-regulated proteins, 20 (26.3%) proteins are specific, 9 (11.8%) proteins are shared with P1GF^−/−^ vs. C57, only one (1.3%) protein is shared with other two groups. Akita. P1GF^−/−^ vs. Akita have 15 specific proteins; 3 proteins are shared with P1GF^−/−^ vs. C57, and one protein is shared with other two groups.

**Figure 7:**
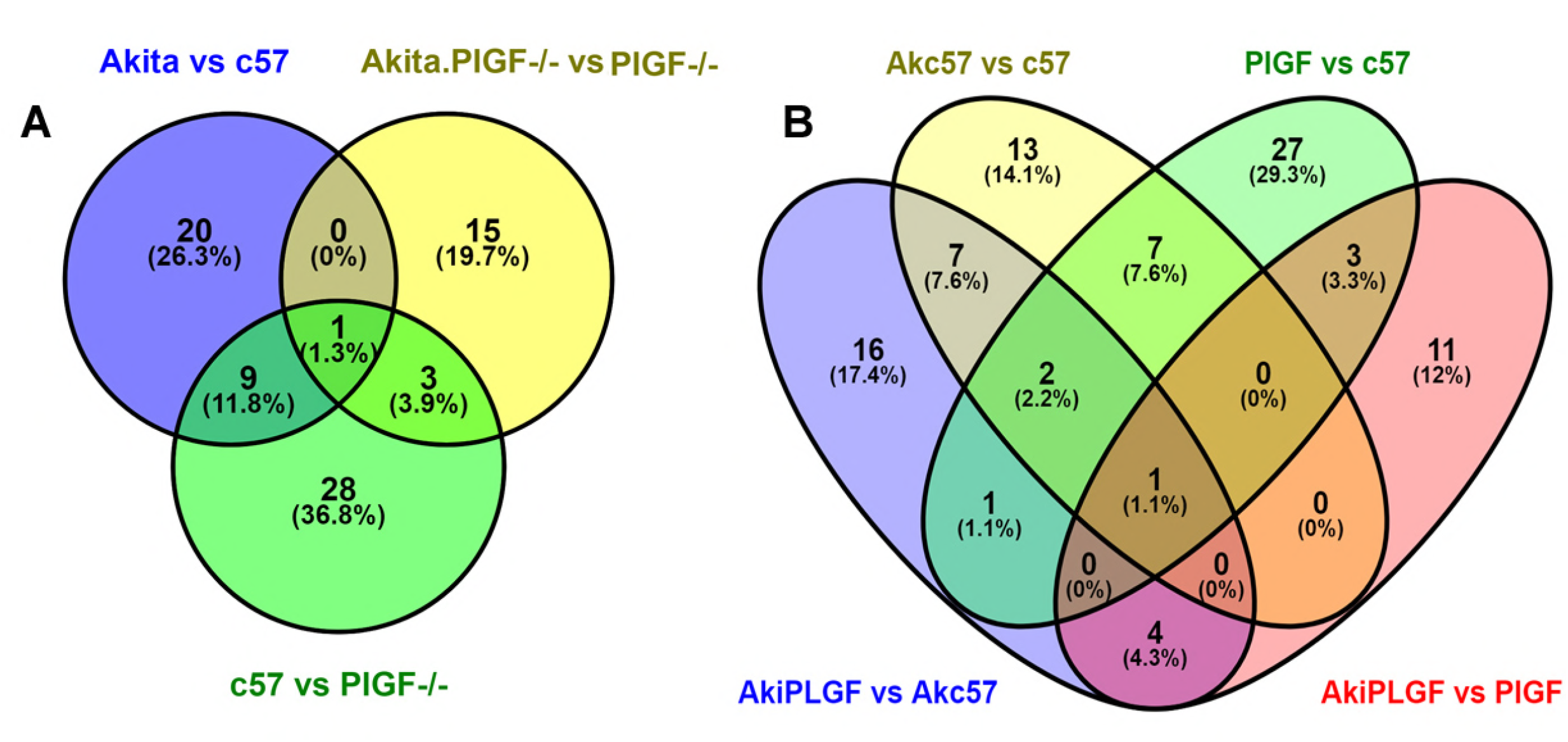
Comparison of each group of up and down-regulated proteins are illustrated by Venn diagrams. (**A**) The three groups of unique up and down-regulated proteins are represented as a Venn diagram. (**B**) The four groups of unique up and down-regulated proteins are represented as a Venn diagram.

## 3.0 Discussion

To elucidate molecular mechanisms that P1GF mediates in early complications in DR, we examine retinal proteome of the four mouse strains Akita.P1GF^−/−^, Akita, C57, and P1GF^−/−^ using label-free mass spec quantification. This approach provides protein information for all retinal cell types during mass spectral analysis. Using normalization by the Z-score method, correlation by Pearson correlation coefficient, we have identified differentially expressed proteins in Akita.P1GF^−/−^ vs Akita (31 proteins), Akita vs C57 (26 proteins), P1GF^−/−^ vs C57 (31 proteins), Akita.P1GF^−/−^ vs. P1GF^−/−^ (19 proteins) combinations, which may be involved in the protective retinal phenotype in diabetes retinopathy conditions.

In Akita.P1GF^−/−^, Map2, Prdx6, Tubb6, Crocc, Hsp90ab1 and Ckb were up-regulated and Gnb2, Snap25, Pcbp1, Atp1a1, Pcbp3, Gnao1, Atp2b1, Gnai2, Vim, Slc25a22, Hnrnpa1, Mecp2 etc., are downregulated when compared to Akita group (**Table 2**). Functional classification and pathway analysis results revealed that Map2 gene encodes microtubule-associated proteins, which are involved in the binding activity of molecular function, cellular component organization or biogenesis, and cellular process. These proteins are thought to be comprised in microtubule assembly, which is a crucial step in neuritogenesis. In rat and mouse, Map2 are neuron-specific cytoskeletal proteins that are augmented in dendrites, associating a role in determining and stabilizing dendritic shape throughout neuron development (Lim & Halpain, 2000). Increased productions of IL-1b, IL-6, or TNF-a decrease the Map2 expression and lead to neuronal cell death. However, the role of the Map2 is not limited to neuronal differentiation or merely neuron cell loss cell marker. Map2 serves to stabilize microtubules (MT) growth by crosslinking microtubules with intermediate filaments. Map2 upregulation increases neuronal cell survival and differentiation. Furthermore Map2 also has a binding domain for the regulatory subunit II of cAMP-dependent protein kinase (PKA). It is known that in diabetes increased oxidative stress and low energy levels contribute to neuron microtubule meshwork degeneration that eventually leads to neuron cell death (Harada et al, 2002; Iriuchijima et al, 2005; Teng et al, 2001). Our results revealed that Map2 protein is up-regulated in Akita.P1GF^−/−^ compared to Akita mice retinas. Peroxiredoxin-6 (Prdx6) protein is up-regulated in the Akita.P1GF^−/−^ compare to Akita and belongs to oxidoreductase class, the antioxidant activity of molecular function, involved in cellular and metabolic process. Prdx isoforms (Prdx) are expressed in the majority of mammalian cells, which differ in structure, catalytic mechanisms and subcellular compartmentalization (Hanschmann et al, 2013). In the retina, Prdx6 was expressed solely by Müller cells and astrocytes. Astrocytes and Müller cells play crucial structural and functional roles in the maintenance of barrier function, uniquely within the retina; decreased Prdx6 is of obvious relevance to any disease where the blood-retinal barrier is compromised, such as diabetic retinopathy, exudative age-related macular degeneration and arterial and venous occlusions (Chidlow et al, 2016).

Tubulin beta6 class V (Tubb6) is a cytoskeletal protein, has binding activity, the structural molecular activity of molecular functions, cellular component organization or biogenesis, cellular process of biological process. Tubb6 allows to modify the organization of muscle microtubules, regardless of the presence or absence of dystrophin, for the guidance and proper organization of microtubules (Oddoux et al, 2013). Crocc (Rootletin) is a class of enzyme modulator, has catalytic activity and is involved in the cellular process. The rootlet aids to anchor the cilium to the cell and functions as a channel for proteins intended for the outer segment of rod photoreceptors (Yang et al, 2005). Hsp90ab1 is a chaperone class of protein that binds to other proteins, thereby stabilizing them in an ATP-dependent manner (Taipale et al, 2010). Creatine kinase B type (Ckb) is a vital enzyme for energy metabolism. It catalyzes the conversion of creatine, consuming adenosine triphosphate (ATP), phosphocreatine and adenosine diphosphate (ADP), in the visual cycle. Ckb is critical for providing energy for the visual cycle in photoreceptors (Zhu et al, 2013). Guanine Nucleotide Binding proteins (Gnb1, Gnb2, Gnb4, Gnai2, Gnao1) are called as G-proteins, which play a key role in insulin signaling pathways. Incessant activation of G-proteins (polymorphism) results in insulin resistance and ultimately increases hepatic glucose output. Gnb gene dimorphism leads to increased cardiac potassium channel activity and increased α-adrenoceptor–mediated vasoconstriction thus resulting in the development and progression of hypertension, obesity and insulin resistance in humans (Benjafield et al, 2001; Poch et al, 2002).

Synaptosomal-associated protein 25 (Snap25) is a component of the trans-SNARE complex, which is proposed to account for the specific membranes fusion and to directly execute fusion by forming a tight complex that brings the synaptic vesicle and plasma membranes together. Zuheng et al. reported that a lowering of Snap25 can be associated with an enhancement of insulin secretion (Ma et al, 2005), but both stimulatory and inhibitory influences by physical interaction of Snap25 with L-Type Ca2+ channels in β cells (Ji et al, 2002). Our mass spectral results are also correlated to Ma et al. (Ma et al, 2005). Snap25 protein is down-regulation in Akita.P1GF^−/−^ group compared to Akita group. Ji et al. supposed to L-Type Ca^2^+ channels interaction with Snap25 and inhibitory and stimulation of insulin, in the same way, our results also supported because of down-regulation of plasma membrane calcium-transporting ATPase1 (Atp2b1 protein) (Ji et al, 2002).

In Akita vs C57 comparison, we have found various classes of up and down-regulated proteins which are involved in the different biological processes, molecular functions, cellular components and Reactome pathways. In particular, we have highlighted Ion transport by P-type ATPases, regulation of insulin secretion and processing of capped intron-containing pre-mRNA. Atp1a1, Atp1b2, and Atp1a2 are up-regulated in Akita compared to the C57 group, which are involved in ROS pathways and lead to obesity, insulin resistance, and metabolic syndrome (Sodhi et al, 2015). Gnb1, Gnb2, and Stxbp1 are involved in regulation of insulin secretion pathway in up-regulation in Akita compared to the C57 group, but these are insulin resistance in up-regulation condition. Andersson et al (Andersson et al, 2012) supported that reduced expression of exocytotic genes (Stxbp1, Snap25, Vamp2) contributes to impaired insulin secretion, and suggested that decreased expression of these genes as part of diabetes condition. Hnrnpk, Hnrnpa1, Hnrnpc, and Hnrnpd are involved in the processing of capped intron-containing pre-mRNA pathway. He and Smith (He & Smith, 2009) suggested that Hnrnp1 and Hnrnp2 are involved in multiple cellular processes, including DNA replication and transcription and mRNA export, as well as pre-mRNA splicing regulation. The upregulation of Hnrnp (Heterogeneous nuclear ribonucleoprotein) proteins on alternative splicing could lead to significant changes in proteomic diversity in a cell population. In our results showed that these Hnrnp proteins are upregulated in Akita compared to C57 but downregulat in Akita.P1GF^−/−^ group.

In P1GF^−/−^ vs C57 group we have identified 31 up and down-regulated proteins, which belong to various classes (**Figure 6**), are involved in different biological process pathways, molecular functions, cellular components and Reactome pathways. Specifically, Gnb1, Gnb2, Gngt1, Gnb4, Vamp2 proteins, which are involved in diabetes directly or indirectly, also up-regulated in P1GF^−/−^ group compared with the C57 group but when compared to Akita are down-regulated.

We have also found up-regulated protein (Ndufa8, Atp5o, Atp5f1) involved in respiratory electron transport, ATP synthesis by chemiosmotic coupling, and heat production by uncoupling proteins. NADH dehydrogenase [ubiquinone] 1 alpha subcomplex subunit 8 (Ndufa8) was mitochondrial intermembrane space assembly pathway. This minor pathway depends on redox-regulated folding events that stabilize, trap and finally import into the mitochondrion the C-X9-C domain-carrying proteins (Schmidt et al, 2010). Eukaryotic translation initiation factor 4α2 (Eif4a2), Eif4a1 are deadenylation of mRNA, upstream of the Eif4a2 gene (Cheyssac et al, 2006), Eif4a1 showed a significant association with type 2 diabetes but which down-regulation in P1GF^−/−^ group compared to the C57 group.

We have also compared Akita to P1GF^−/−^, significantly Hexokinase (Hk1), Stxbp1, Hnrnpc, Vamp2, Vamp3, Atp1b1, Pcbp1, Pcbp2, Slc6a11, and Nono are up-regulated in Akita, which is mostly involved in diabetes. Rcvrn (Recoverin), Sag, Epb4, Crocc, Cplx4, Gnat1, and Gngt1 are down-regulated in Akita group. Recoverin protein is a neuronal calcium-binding protein that is primarily detected in the photoreceptor cells of the eye. It plays a crucial role in the inhibition of rhodopsin kinase, a molecule which regulates the phosphorylation of rhodopsin. A reduction in this inhibition helps regulate sensory adaptation in the retina, since the light-dependent channel closure in photoreceptors causes calcium levels to decrease, which relieves the inhibition of rhodopsin kinase by calcium-bound recoverin (Chen et al, 1995).

Our results also correlated the earlier studies because the recoverin is known to be down regulated in diabetic condition. Akita.P1GF^−/−^ vs. P1GF^−/−^ combination, significantly Tubb3, Tubb6, Tubb2b, Tubb5, Hspa8, Hsp90ab1 were up-regulated while Atp5o, Map1b, Hnrnpm, Vdac2, Pura, Cox5b Ndufa8 and Erh proteins are down-regulated (**Table 4**) in Akita.P1GF^−/−^ compared to P1GF^−/−^.

## 4.0 Conclusion

Here we have analyzed the proteomics changes in associated with P1GF ablation in diabetic and healthy condition. Identifying the number of the proteins that can contribute to better understanding molecular mechanisms of P1GF function its target and off-target effects.

Reduced insulin resistance of Akita.P1GF^−/−^ are potentially associated with P-type ATPases and GNB group proteins involve in cell metabolism that allow retinal cells to better utilize the glucose reducing the metabolic stress and ROS production. Particularly major neuron survival factor MAP2 and antioxidant defence protein Prdx6 are upregulated in P1GF^−/−^ and Akita.P1GF^−/−^ conditions. Increased expression of Tubb and Hsp protein groups in Akita.P1GF^−/−^ animals indicated increase cell chaperon activity and integrity that potentially increases retinal cell survival. We intend this proteome wide analysis will encourage further research into identified pathways of P1GF effects in DR.

## 5.0 Materials and Methods

### 5.1 Mouse Strains

The use of animals was in compliance with the Association for Research in Vision and Ophthalmology (ARVO) Statement for the Use of Animals in Ophthalmic and Vision Research and approved by the Institutional Animal Care and Use Committee of The Johns Hopkins University (Protocol number: M016M480). Mice were housed at the special pathogen-free (SPF) Cancer Research Building Animal Facilities at Johns Hopkins Hospital. Mouse strains

C57BL/6-Ins2<Akita>/J.P1GF^−/−^, C57BL/6-P1GF^−/−^, C57BL/6-Ins2<Akita>/J, and C57BL/6J (C57) were generated and maintained as described previously. (Huang et al, 2015b) Briefly, Akita mice were crossed with P1GF^−/−^ mice (Van de Veire et al, 2010) in a C57BL/6J background for two generations to give birth to the progeny with the genotype of Akita.P1GF^−/−^. The breeding program graphical abstract is presented in the **Figure 1A**.

## 5.3 Verification of diabetic conditions

Diabetic phenotype and genotype were confirmed 4.5 weeks after birth by blood glucose >250 mg/dL (One-Touch Lifescan meter; Lifescan, Inc., Milpitas, CA) in a drop of blood from a tail puncture Diabetes development were not maintained with insulin. **Figure 1B** The levels of blood glucose were converted to glycated hemoglobin (HbA1c) using an online tool (https://www.accu-chek.com/us/glucose-monitoring/a1c-calculator.html). Final blood glucose concentration and body weight were measured before sacrifice.

## 5.4 Retinal samples collection

Mice were euthanized by ketamine overdose (100 mg/kg), intraperitoneal injection. Mouse eyes were enucleated, and retinas were carefully isolated free of the cornea, lens, vitreous humor and retinal pigment epithelium (RPE) on ice in cold phosphate-buffered saline (PBS, 0.1M, pH=7.2) under the dissection microscope Stemi 2000-C (Zeiss). Retinas were frozen with liquid nitrogen and then stored at −80°C until further use. (**Figure 1C**)

## 5.5 Protein extraction and peptide preparation

Retinas were homogenized using RIPA buffer (R0278-50ML; Sigma) supplemented with Protease and Phosphates Inhibitor cocktail (Cell Signalling Technology). Retinal homogenates were centrifuged (12,000g, 4°C, and 10min) to remove tissue debris. The protein concentration of the supernatant was determined using Pierce BCA Protein Assay Kit (Thermo Fisher Scientific, Cat#: 23225); 50-microgram of total protein from each sample was used for peptide preparation. DTT reduced proteins and alkylated by iodoacetamide (10mM, in 50mM Tris-HCl, pH8.0) at room temperature for 1h. The protein samples were then digested with trypsin/LysC (Promega, Cat#: V5073) in 25mM ammonium bicarbonate solution at 37°C for 12 h. The resultant peptides were washed and desalted with the desalting column (Pierce Spin-Tip) and eluted with 50μl 5% acetonitrile + 0.4% trifluoroacetic acid. The peptides mixtures were run through the ziptip C18 (Millipore) and then dried with SpeedVac concentrator. (**Figure 1D**)

## 5.3 Mass spectrometry

The LC/MS/MS analysis of samples was carried out using a Thermo Scientific Q-Exactive hybrid Quadrupole-Orbitrap Mass Spectrometer and a Thermo Dionex UltiMate 3000 RSLCnano System (Poolchon Scientific, Frederick, MD, USA). Peptide mixtures from each sample were loaded onto a peptide trap cartridge at a flow rate of 5 μL/min. The trapped peptides were eluted onto a reversed-phase PicoFrit column (New Objective, Woburn, MA) using a linear gradient of acetonitrile (3-36%) in 0.1% formic acid. The elution duration was 120 min at a flow rate of 0.3 μl/min. Eluted peptides from the PicoFrit column were ionized and sprayed into the mass spectrometer, using a Nanospray Flex Ion Source ES071 (Thermo) under the following settings: spray voltage, 1.6 kV, Capillary temperature, 250°C.

## 5.4 Label-free proteomics data analysis

The mass spectral (MS) raw data were analyzed with MaxQuant computational proteomics platform (version 1.6.1.0) and its built-in Andromeda search engine (Cox & Mann, 2008). The LTQ-Orbitrap peptides were identified (main search peptide search = 4.5 ppm and 20 ppm for first search peptide tolerance, respectively) from the MS/MS spectra searched against *Mus musculus* UniProtKB (83,598 entries) target database (Perumal et al, 2014) using Andromeda search engine. (Cox et al, 2011) This target database was also shared with the common contaminants and concatenated with the reversed versions of all sequences. In group-specific parameters, enzyme-specific was set to trypsin, and two missed cleavages were allowable; variable modifications were selected as fixed modification whereas carbamidomethylation (C) and oxidization (M) were set as fixed and variable modifications respectively were selected as variable modifications and 5 set to a maximum number of modifications per peptide. The type was set the standard; multiplicity was set to 1 to account for the label-free state, and Label-free quantification (LFQ) was set LFQ, 2 set for minimum ration count (Cox & Mann, 2008). The FDR (false discovery rate) was set to 0.01 for proteins, peptides and other parameters were set to default. Label minimum ratio count was set to 2, peptides for quantification was set to unique and razor and re-quantify to allow identification and quantification of proteins in groups for LFQ analysis.

First, MaxQuant corrects for systematic inaccuracies of measured peptide masses and corresponding retention times of extracted peptides from the raw data. Then for peptide identification, mass and intensity of the peptide peaks in mass spectrometry (MS) spectrum are detected and assembled into three-dimensional (3D) peak hills over the m/z retention time plane, which is filtered by applying graph theory algorithms to identify isotope patterns. High mass accuracy is achieved by weighted averaging and through mass recalibration by subtracting the determined systematic mass error from the measured mass of each MS isotope pattern. Peptide and fragment masses (in case of an MS/MS spectrum) are searched in an organism-specific sequence database and are then scored by a probability-based approach termed peptide score. For limiting a certain number of peak matches by chance, a target-decoy-based false discovery rate (FDR) approach is utilized. The FDR is determined using statistical methods that account for multiple hypotheses testing. Also, the organism-specific database search includes not only the target sequences but also their reverse counterparts and contaminants, which helps to determine a statistical cut-off for acceptable spectral matches. The assembly of peptide hits into protein hits to identify proteins is the next step, in which each identified peptide of a protein contributes to the overall identification accuracy. Also, an FDR-controlled algorithm called matching between runs is incorporated, which enables the MS/MS free identification of MS features in the complete data set for every single measurement, leading to an increase in the number of quantified proteins per sample (Cox & Mann, 2008).

## 5.5 Perseus analysis pipeline

The MaxQuant software generated output file “proteingroups.txt” was utilized for Pearson correlation, clustering and statistical analysis using Perseus software version 1.6.1.1 (Tyanova et al, 2016). Unverified hierarchical clustering of the LFQ values was carried out based on Euclidean distances on the Z-scored among mean values. For statistical analysis, two-samples t-test-based statistics with P < 0.05 was applied on Log2 transformed LFQ values and the minimum number of values “in at least one group” is 3 to assert proteins regulation as significant for the specific groups (Cox et al, 2014; Perumal et al, 2015).

## 5.6 Functional classification and pathway analysis

Proteins determined to be differentially expressed as illustrated based on the data in our LFQ experiments were tabularize in Excel and their gene names were used for functional annotation and pathways analysis. First, DAVID tool (v6.7) (Huang da et al, 2009) (http://david.abcc.ncifcrf.gov/home.jsp) was used for interpreting the GOBP terms of the differentially-expressed proteins. The protein list was uploaded into DAVID and searched for enrichment for GOBP term, and the results were filtered based on threshold count ≥ 2 and P values < 0.05.

## 5.7 Protein-protein network analysis

We have also performed the functional enrichment and interaction network analysis using STRING 10.5 database (Szklarczyk et al, 2011) on differentially expressed proteins. SPRING tool classified the proteins according to the Gene Ontology (GO) categories such as biological process (BP), molecular function (MF), Cellular Component (CC) and KEGG (Kyoto Encyclopedia of Genes and Genomes database) pathways. (Kanehisa et al, 2017) The tool Venny 2.1 (http://bioinfogp.cnb.csic.es/tools/venny/) was used to generate Venn diagrams.

## 5.9 Statistical analysis

All values were expressed as the mean ± standard deviation (SD) for the respective groups. Statistical analyses were performed with GraphPad Prism software (https://www.graphpad.com/scientific-software/prism/). The Student’s t-test, one-way ANOVA test with Tukey multiple comparisons, were used. P value less than 0.05 were considered significant.

## 5.10 Accession numbers

The accession number for the raw mass spectrometry data reported in this study is ProteomeXchange Consortium: (*Mass Spec Raw data will be uploaded during the revision process*)

---

**Supplementary Figure 1:** Quantitative profiling of retinal tissue between Akita.P1GF^−/−^ vs. Akita groups. **A.** Hierarchical clustering of Z-scored median LFQ intensities for all proteins in Akita.P1GF−/− vs. Akita groups. **B.** LFQ intensity of each protein across Akita.P1GF−/− vs. Akita group samples. **C.** LFQ intensity of each protein across Akita.P1GF−/− vs. Akita group samples after imputation. **D.** Pearson correlation coefficient of LFQ intensities among LC-MS/MS runs in Akita.P1GF−/− vs. Akita groups.

**Supplementary Figure 2:** Quantitative profiling of retinal tissue between Akita vs. C57 groups. **A.** Hierarchical clustering of Z-scored median LFQ intensities for all proteins in Akita vs. C57 groups. **B.** LFQ intensity of each protein across Akita vs. C57 group samples. **C.** LFQ intensity of each protein across Akita vs. C57 group samples after imputation. **D.** Pearson correlation coefficient of LFQ intensities among LC-MS/MS runs in Akita vs. C57 groups.

**Supplementary Figure 3:** Quantitative profiling of retinal tissue between P1GF^−/−^ vs. C57 groups. **A.** Hierarchical clustering of Z-scored median LFQ intensities for all proteins in P1GF^−/−^ vs. C57 groups. **B.** LFQ intensity of each protein across P1GF^−/−^ vs. C57 group samples. **C.** LFQ intensity of each protein across P1GF^−/−^ vs. C57 group samples after imputation. **D.** Pearson correlation coefficient of LFQ intensities among LC-MS/MS runs in P1GF^−/−^ vs. C57 groups.

**Supplementary Figure 4:** Quantitative profiling of retinal tissue between Akita.P1GF^−/−^ vs P1GF^−/−^ groups. **A.** Hierarchical clustering of Z-scored median LFQ intensities for all proteins in Akita.P1GF−/− vs P1GF^−/−^ groups. **B.** LFQ intensity of each protein across Akita.P1GF−/− vs P1GF^−/−^ group samples. **C.** LFQ intensity of each protein across Akita.P1GF−/− vs. P1GF^−/−^ group samples after imputation. **D.** Pearson correlation coefficient of LFQ intensities among LC-MS/MS runs in Akita.P1GF−/− vs P1GF^−/−^ groups.

**Supplementary Figure 5:** Representative mass spectra of **A.** Gnb1, **B.** Gnb2, **C.** Prdx6 and **D.** Map2 proteins.

## Acknowledgments

The authors wish to acknowledge the contribution of: Lijuan Fan for technical assistance, Dmitry Rumyancev for artwork design, and Jianjiang Hao for MS analysis.

**Authors Contributions**
The study was conceived and designed by M.S.S., A.L. and H.H. H.H. performed the animal handling, sample collection and *in vivo* examinations. The manuscript was written by M.S.S, A.L.,H.H. and critically revised by H.H. and S.T. All Authors reviewed and accepted the final version of the manuscript.

